# Apomorphine susceptibility and prenatal infection alter neurodevelopment, synaptic density and anticipatory behavior in rats

**DOI:** 10.1101/2025.08.25.672101

**Authors:** Kate Witt, Meyrick Kidwell, Janine Doorduin, Erik. F.J. de Vries, Iris E. Sommer, Darren J. Day, Bart A. Ellenbroek, Cyprien G.J. Guerrin

**Affiliations:** Department of Nuclear Medicine and Molecular Imaging, University Medical Center Groningen, University of Groningen, Hanzeplein 1, 9713 GZ, Groningen, the Netherlands; Behavioural Neurogenetics Group, Victoria University of Wellington, PO Box 600, Wellington, New Zealand; Department of Biomedical Sciences, University Medical Center Groningen, University of Groningen, 9713 GZ Groningen, The Netherlands; School of Biological Sciences, Victoria University of Wellington, PO Box 600, 6104, Wellington, New Zealand; Department of Medical Neuroscience, Donders Institute for Brain, Cognition and Behavior, Radboud University Medical Center, Nijmegen, the Netherlands

**Keywords:** Schizophrenia, Maternal immune activation, APO-SUS, polygenic susceptibility, rat behavior

## Abstract

Schizophrenia is a complex psychiatric disorder, driven by genetic and environmental factors. While individual risk genes have limited impact, polygenic susceptibility increases the likelihood of schizophrenia and heightens sensitivity to environmental stressors, such as prenatal immune activation. Yet, preclinical studies often focused on single-gene mutations, leaving polygenic influences largely unexplored. Using the apomorphine-susceptible (APO-SUS) rat model, which exhibits schizophrenia-like features, we investigated how polygenic susceptibility influences early neurodevelopment, synaptic density, and behavior, and how these effects are modulated by prenatal immune activation. APO-SUS rats demonstrated early neurodevelopmental abnormalities, including a reduced number and duration of separation-induced ultrasonic vocalizations (USVs), increased principal frequency of USVs, and reduced heart rate variability (HRV), indicative of heightened sympathetic dominance commonly seen in psychiatric disorders. These effects were particularly pronounced in females. Male APO-SUS rats exhibited elevated synaptophysin levels, a presynaptic marker for synaptic density, in the frontal cortex during adolescence and in the hippocampus during adulthood. Interestingly, prenatal immune activation counteracted some of these changes, preventing HRV reduction and normalizing synaptophysin levels. Male and female APO-SUS rats, as well as Wistar male rats exposed to prenatal immune activation, showed anticipatory behavior during adolescence, but not in adulthood. Our results suggest that polygenic susceptibility induces early neurodevelopmental changes and that genetic and environmental risk factors do not always act synergistically; sometimes counterbalancing each other. Future studies should explore how early neurodevelopmental changes, such as alterations in USVs and HRV, influence later behavioral outcomes in polygenic models of schizophrenia.

**Highlights:**

- Apomorphine-susceptible rats show altered vocalizations and heart rate variability
- Apomorphine susceptibility increases synaptophysin in frontal cortex and hippocampus
- Prenatal immune activation reduces synaptophysin in the adult frontal cortex
- Prenatal immune activation prevents apomorphine-susceptible brain and behavior changes

## 1. Introduction

Schizophrenia is a major psychiatric disorder affecting approximately 1% of the world’s population (Tandon et al., 2013). Its symptomatology includes positive (hallucinations, delusions), negative (anhedonia, apathy, amotivation), mood, and cognitive symptoms (Tandon et al., 2013). Schizophrenia affects both sexes, with differences in onset, clinical course and treatment response (Sommer et al., 2020). Understanding the etiology of schizophrenia is challenging due to its multifactorial nature. Genetic factors play a significant role as highlighted by twin, adoption, and family studies (Riley and Kendler, 2006). However, genome-wide association studies reveal that individual genes contribute minimally to the overall risk (Alnæs et al., 2019; Gasse et al., 2019; Ripke et al., 2014). Instead, the cumulative effect of multiple risk genes, summarized as polygenic risk scores, significantly increases the likelihood of specific schizophrenia-related traits, such as treatment resistance and cortical thinning (Alnæs et al., 2019; Gasse et al., 2019; Ripke et al., 2014). Additionally, environmental influences, such as prenatal infections, childhood trauma and growing up in large cities have been associated with an increased risk of developing schizophrenia and other psychiatric disorders, including autism spectrum disorder, depression, and bipolar disorder (Modai and Shomron, 2016; Uher, 2014).

Schizophrenia likely arises from altered neurodevelopmental processes—including neurogenesis, neuronal differentiation, synaptogenesis, and synaptic pruning—during key developmental windows (Cardozo et al., 2019; Germann et al., 2021; Paolicelli et al., 2011). Disruptions in these processes may lead to reduced synaptic density in later life. For instance, positron emission tomography (PET) imaging and post-mortem studies have revealed reductions in synaptic density markers, such as synaptic vesicle glycoprotein 2A (SV2A), synaptophysin, synaptosomal-associated protein 25 (SNAP25), and postsynaptic density protein 95 (PSD95), in the frontal cortex and hippocampus, which are areas critical for cognitive and emotional processing in individuals with schizophrenia (Corradini et al., 2009; Onwordi et al., 2020; Osimo et al., 2019). However, the extent to which polygenic risk scores and environmental factors, such as prenatal infection, contribute to these neurodevelopmental alterations remains unclear. While prospective studies in patients present significant challenges, these interactions can be modeled in rodents, offering valuable insights into the disorder’s pathophysiology.

Previous research focused on single-gene mutations combined with environmental stressors, yet found limited effects, leaving polygenic susceptibility largely unexplored (Guerrin et al., 2021). Given the challenges of modeling polygenic risk with multiple engineered mutations, selectively bred rodent lines like the apomorphine-susceptible (APO-SUS) rats offer a practical alternative (Cools et al., 1993). Bred for heightened sensitivity to the dopamine agonist apomorphine, APO-SUS rats display increased dopaminergic tone—an important feature of schizophrenia pathophysiology (Cools et al., 1990; Ellenbroek et al., 1995; Oliveras et al., 2023). APO-SUS rats show schizophrenia-relevant phenotypes, including reduced prepulse and latent inhibition, anhedonia (reduced sucrose preference), increased sensitivity to stimulants, hyperactivity, and mild cognitive deficits (Cools et al., 1993; Ellenbroek et al., 1995; Ellenbroek and Cools, 2000; Maas et al., 2020; van der Elst et al., 2006; van Vugt et al., 2014). Neurobiologically, they exhibit elevated dopamine D2 receptor binding, hypothalamic-pituitary-adrenocortical axis hyperactivity, hypomyelinated GABAergic interneurons, oxidative stress, mitochondrial dysfunction, and immune alterations (e.g., reduced T/B cell ratios, decreased NK cells) (Ellenbroek and Cools, 2002; Maas et al., 2020; Rots et al., 1996). These abnormalities mirror those found in schizophrenia, including presynaptic dopamine elevation, myelin dysregulation, GABAergic deficits, immune dysregulation, and oxidative stress (Doorduin et al., 2009; Du et al., 2013; Fillman et al., 2016; Flynn et al., 2003; Glausier and Lewis, 2017; Hakak et al., 2001; Howes and Murray, 2014; Ng et al., 2008; Rajasekaran et al., 2015; Schmidt and Mirnics, 2014; Selten et al., 2016). Furthermore, GWAS meta-analyses have identified over 100 schizophrenia-associated loci, implicating genes related to dopaminergic signaling (e.g., *DRD2*), glutamatergic transmission (*GRIN2A, GRM3, GRIA1*), calcium channels, and immune function (Ripke et al., 2014; Sekar et al., 2016). In the APO-SUS rat model, a cluster of genetic variations including topoisomerase II-based recombination hotspots, and region-specific epigenetic modifications such as DNA methylation differences in the cerebellum and hypothalamus contribute to a complex phenotype resembling schizophrenia, affecting neural development, neuronal signaling, and gene expression regulation (Coolen et al., 2005; van Loo and Martens, 2007; Van Schijndel et al., 2010). APO-SUS rats also exhibit alterations in *KCNIP1* and *APH1B*, involved in potassium channel regulation and γ-secretase signaling (Coolen et al., 2005; van Loo and Martens, 2007; Van Schijndel et al., 2010). Related genes (*KCNJ13, KCNV1, APH1A*) have been identified in schizophrenia GWAS (Ripke et al., 2014). Despite belonging to distinct subfamilies, they converge on key pathways—neuronal excitability and neurodevelopmental signaling—further supporting the face validity of APO-SUS rats as an idiopathic, polygenic model of schizophrenia vulnerability (Coolen et al., 2006, 2005; Ellenbroek and Karl, 2016)

In this study, we investigated the impact of polygenic apomorphine susceptibility in APO-SUS rat, prenatal exposure to infection, and their combined effect on neurodevelopment, behavior, and synaptic density. To model maternal infection, we treated pregnant dams with the viral mimic polyinosinic:polycytidylic acid (poly I:C), which induces a range of behavioral and neuronal abnormalities in the offspring that are relevant to several neurodevelopmental disorders, including schizophrenia and autism (Brown and Meyer, 2018; Careaga et al., 2017), and has shown varied outcomes, sometimes protective or synergistically exacerbating symptoms (Guerrin et al., 2021). On postnatal days (PND) 7 and 14, we assessed separation-induced ultrasonic vocalizations (USVs) as a marker of social-emotional communication and early neurodevelopment (Brudzynski, 2013; Shair, 2007; Wöhr and Schwarting, 2013), alongside heart rate variability (HRV), a measure of autonomic function that is associated with future health outcomes and the severity of neuropsychiatric disorders, such as schizophrenia, autism, and bipolar disorders (Benjamin et al., 2021; Haigh et al., 2021; Kidwell and Ellenbroek, 2018; Rajendra Acharya et al., 2006). To explore changes in anticipatory apathy—a domain often disrupted in schizophrenia—we evaluated anticipatory behavior during adolescence and adulthood (Gard et al., 2007; Sherdell et al., 2012). Synaptic integrity and density were examined via western blot analyses of synaptophysin and postsynaptic density protein 95 (PSD95) in the hippocampus and prefrontal cortex, providing insight into the neurobiological underpinnings of schizophrenia-related behaviors (Barch and Ceaser, 2012; Coley and Gao, 2018; Wiedenmann et al., 1986). Given the sex-dimorphic nature of schizophrenia, we included both male and female rats.

## 2. Materials and Methods

All detailed protocols, including specific experimental setups, data acquisition workflows, and reagent preparations, are provided in the Supplementary Materials.

### 2.1. Animals

All experiments were approved by the Victoria University of Wellington’s Animal Ethics Committee and conducted in accordance with its animal care principles. Male and female Wistar and APO-SUS rats were used. APO-SUS rats, Wistar rats selectively bred for apomorphine susceptibility (>500 gnaws in 45 min after 1.5 mg/kg apomorphine injection), exhibit schizophrenia-relevant behavioral alterations independent of apomorphine.

Pregnant dams were housed individually after mating and checked daily for vaginal plugs (gestational day 1). Offspring were weaned on postnatal day (PND) 21 and housed in same-sex groups of 2–5 rats. Animals were maintained under controlled conditions (21 ± 2 °C, 55–60% humidity, 12:12-h reversed light cycle) with ad libitum food and water.

### 2.2. Experimental Design

Male and female offspring were divided into four experimental groups: (1) control (offspring of saline-treated Wistar dams), (2) MIA (offspring of poly I:C-treated Wistar dams), (3) APO-SUS (offspring of saline-treated APO-SUS dams), and (4) APO-SUS+MIA (offspring of poly I:C-treated APO-SUS dams).

Behavioral and physiological assessments were performed at three developmental stages: PND7 and PND14 (USVs and HRV), adolescence (PND31–40; anticipatory locomotor activity), and adulthood (PND56–66; anticipatory locomotor activity). Different cohorts were used for early life, adolescence and adulthood to avoid repeated testing effects and to collect the brains for Western Blot analysis.

### 2.3. Maternal Immune Activation

Maternal immune activation (MIA) was induced by injecting pregnant rats subcutaneously with polyinosinic:polycytidylic acid (poly I:C, 5 mg/kg body weight) or saline on gestational day 15. This time point, commonly used in rat MIA models, corresponds to a critical period of cortical neuron migration, neurogenesis, synaptogenesis, myelination, and microglial colonization in the fetal rat brain (Guerrin et al., 2022, Guerrin et al., 2021; Murray et al., 2019; Sarkar et al., 2019). GD15 also approximates the early second trimester of human pregnancy, a developmental window particularly associated with increased risk for schizophrenia, as well as emotional, social, and attentional disturbances in offspring (Hardie et al., 2025). Maternal weight and body temperature were recorded before and 2 and 24 hours after the injection to confirm immune activation. Treated dams were returned to their cages and left undisturbed for the remainder of pregnancy.

### 2.4. Electrocardiographic data and USVs

HRV was assessed on PND7 and PND14 via electrocardiograms (ECG) using ECGenie. Pups were placed individually in a temperature-controlled ECG holder, and heartbeats were recorded for 8 minutes via dedicated paw electrodes and analyzed using LabChart 8 software. During the ECG recordings, USVs were captured using an ultrasound-sensitive microphone (ultraMic 250 kHz) placed above the ECG holder and analyzed with DeepSqueak software.

### 2.5. Anticipatory Locomotor activity

Anticipatory locomotor activity was assessed in adolescence (PND31–40) and adulthood (PND56–66) using activity chambers (42×42×30 cm) equipped with infrared sensors. The experiment lasted 10 days, divided into habituation (days 1–4), pre-anticipation (days 5–6), learning (day 7-8) and anticipation (days 9–10) phases. Distance traveled and the number of rearings were measured during the first 25 minutes of each session. Rewards (Froot Loops/ sugary cereals) were introduced in the chambers after 25 minutes during the learning and anticipation phases.

### 2.6. Western Blot Analysis

Protein levels of synaptophysin and PSD95 were assessed in the hippocampus and frontal cortex of adolescent and adult rats. After preparation of the brain samples, total protein concentrations were measured, then proteins were separated and probed with antibodies for synaptophysin and PSD95. Signals were visualized using a Typhoon scanner, and band intensities were analyzed with Fiji (ImageJ) software. Regions of interest were drawn around the target bands, and background intensity was subtracted to calculate corrected band intensities, which were normalized to the median intensity of the corresponding gel.

### 2.7. Statistical Analysis

Statistical analyses for USV, HRV, and locomotion were conducted using SPSS (IBM SPSS Statistics, Version 22.0). Generalized Estimating Equations (GEE) were applied to these longitudinal datasets, incorporating Genotype, MIA, and Time as fixed factors. GEE was selected over repeated-measures ANOVA because it accounts for within-subject correlations, accommodates unequal group sizes, allows for the inclusion of missing data without listwise deletion, and does not require the assumption of sphericity. These features made GEE particularly suitable for our dataset, which involved a limited number of repeated measures per animal, occasional missing data points, and modest to small sample sizes in some experimental conditions. Post-hoc pairwise comparisons following GEE were conducted using the Least Significant Difference (LSD) procedure, corrected for multiple comparisons. Given the established sex differences in psychiatric disorders, all statistical analyses were performed separately for males and females. This approach allowed us to account for potential sex-specific effects while maintaining statistical power, particularly in light of small group sizes in certain conditions. Western blot data, which did not involve repeated measures, were analyzed using two-way ANOVAs with Genotype and MIA as between-subject factors. Western blot data analyses were conducted separately for sex (male/female), brain region (frontal cortex/hippocampus), and age (adolescence/adulthood). Tukey’s post-hoc tests were used for ANOVA analysis. Prior to parametric testing, data were assessed for normality using the Shapiro-Wilk test and for homogeneity of variance using Levene’s test.

Outliers were screened using boxplots and z-scores (|z| > 3.29) but were retained in the analysis unless attributable to clear experimental error. Due to errors in sex determination at PND7, four pups initially classified as males (two in the MIA group and two in the APO-SUS+MIA group) were later reassigned to the female group. This resulted in a reduced sample size for males in these conditions (n reduced by two) and a corresponding increase in the female groups. In addition, technical issues such as excessive movement during HRV recordings or signal acquisition failure led to data loss in five animals: one female control and four female MIA pups on PND7. In the male group, one MIA animal lacked usable HRV data on PND14. These issues resulted in some final group sizes as low as n = 3 or 4 in certain timepoints for MIA males in HRV and USV measures.

Full statistical results, including Wald Chi-square values and degrees of freedom for the GEE analyses, F-values and degrees of freedom for the two-way ANOVAs, and p-values for all main effects and interactions, are reported in Supplementary Table 1. For clarity, direct comparisons between MIA and APO-SUS groups were not included, as these groups differ on two independent variables. Data are presented as mean ± standard deviation (SD).

The main effects of “Genotype,” “MIA,” and “Time,” along with their interactions, are summarized in Supplementary Table 1 (Appendix), including statistical details from GEE and Two-Way ANOVA (*W, F, df*, and *p*-values).

## 3. Results

### 3.1. APO-SUS rats had a reduced average USV call length and increased principal frequency

The following USV parameters were measured: number of calls, average length of the calls, total length of the calls, and principal frequency (Figure 1).

**Figure 1.**
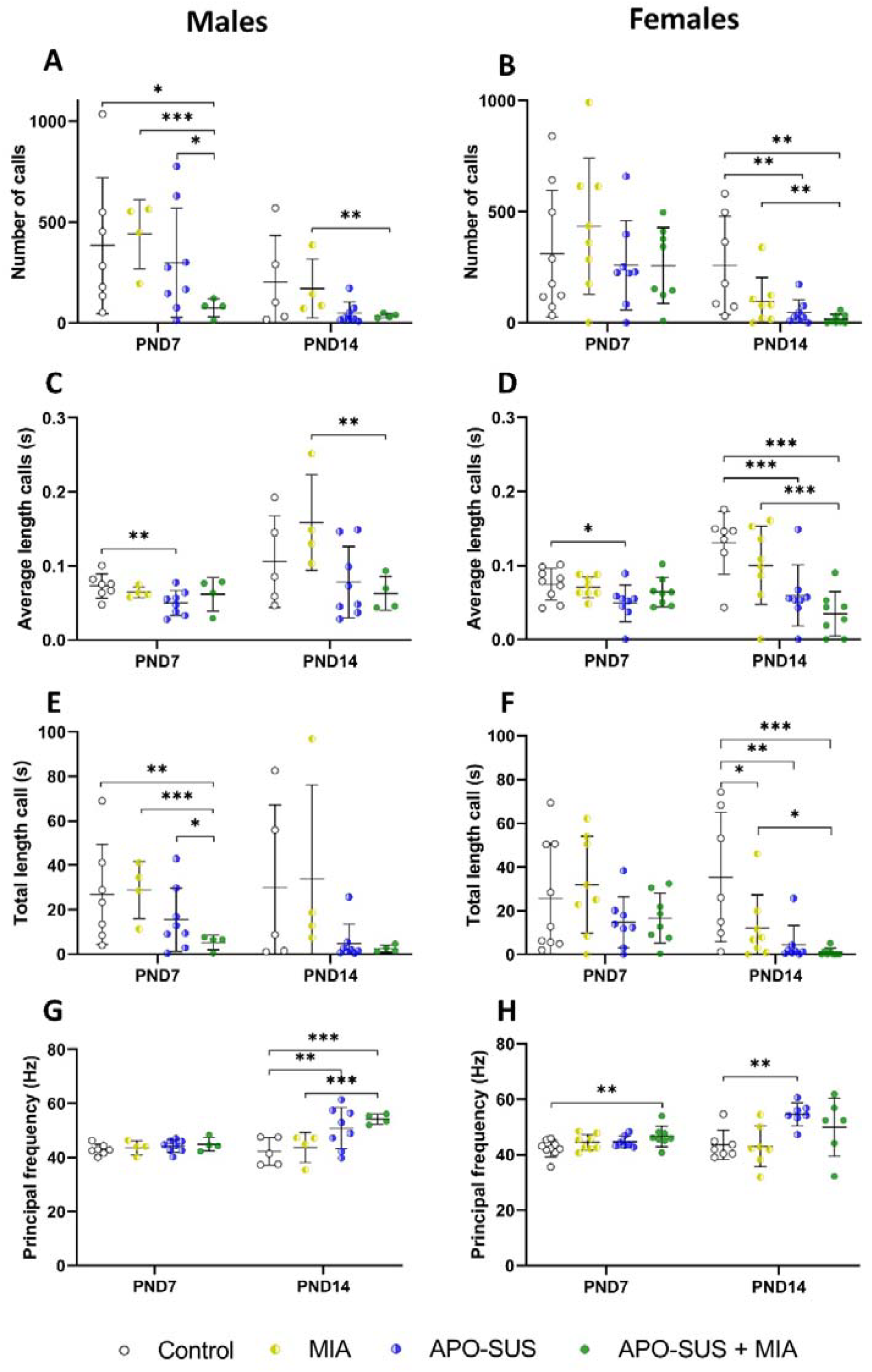
Ultrasonic vocalizations in control, MIA, APO-SUS, and APO-SUS+MIA rats. Number of calls in males (**A**.) and females (**B**.). Average length of the calls (s) in males (**C**.) and females (**D**.). Total length of the call (s) in males (**E**.) and females (**F**.). Principal frequency (Hz) in males (**G**.) and females (**H**.). N=4-8 males and N=8-9 females per group. Data is presented as mean± SD. Statistically significant differences between groups are indicated by asterisks: *p<0.05, **p<0.01, ***p<0.001. Significant differences between time points and between MIA and APO-SUS are not shown.

In males, GEE analysis revealed a main effect of Genotype on all USV measures (p < 0.001), and Time × Genotype × MIA interactions on average length calls (p=0.035). Post-hoc tests indicated that on PND7, APO-SUS+MIA rats emitted fewer calls and had a shorter total call length than control (p < 0.01), MIA (p < 0.001), and APO-SUS rats (p < 0.05). APO-SUS rats also had a shorter average call length than controls on PND7 (p < 0.01). On PND14, APO-SUS+MIA rats emitted fewer calls than MIA rats (p < 0.05) and had a shorter average call length than MIA rats (p < 0.001). APO-SUS rats had a higher principal frequency than controls on PND14 (p < 0.01), and APO-SUS+MIA rats had a higher frequency than both control and MIA rats (p < 0.001).

In females, GEE analysis showed a main effect of Genotype on all USV parameters (p < 0.01), a Genotype x Time interaction on average length calls (p < 0.001), and principal frequency (p = 0.023), and a MIA x Time interaction on average length calls (p = 0.020). Post-hoc comparisons revealed that on PND14, APO-SUS and APO-SUS+MIA rats emitted fewer calls than controls (p < 0.01), and APO-SUS+MIA rats also emitted fewer calls than MIA rats (p < 0.05). APO-SUS rats had a shorter average call length than controls on both PND7 and PND14 (p < 0.05), while APO-SUS+MIA rats had shorter average call length than both controls and MIA rats on PND14 (p < 0.001). Total call length was reduced in APO-SUS, APO-SUS+MIA, and MIA rats compared to controls (p < 0.05), and was also lower in APO-SUS+MIA rats than in MIA rats (p < 0.05). Finally, APO-SUS+MIA rats showed a higher principal frequency than controls on PND7 (p < 0.01), and APO-SUS rats had a higher principal frequency than controls on PND14 (p < 0.001).

### 2.3. APO-SUS rats had a reduced heart rate and heart rate variability

Heart rate variability was measured using the average heart rate (Figure 2.A.B.), the low frequency (Figure 2.C.D.), the high frequency (figure 2.E.F.), the low frequency/high frequency (LF/HF) ratio (Figure 2.G.H.), and the root mean square of successive heartbeat internal differences (RMSSD) (figure 2.I.J.).

**Figure 2.**
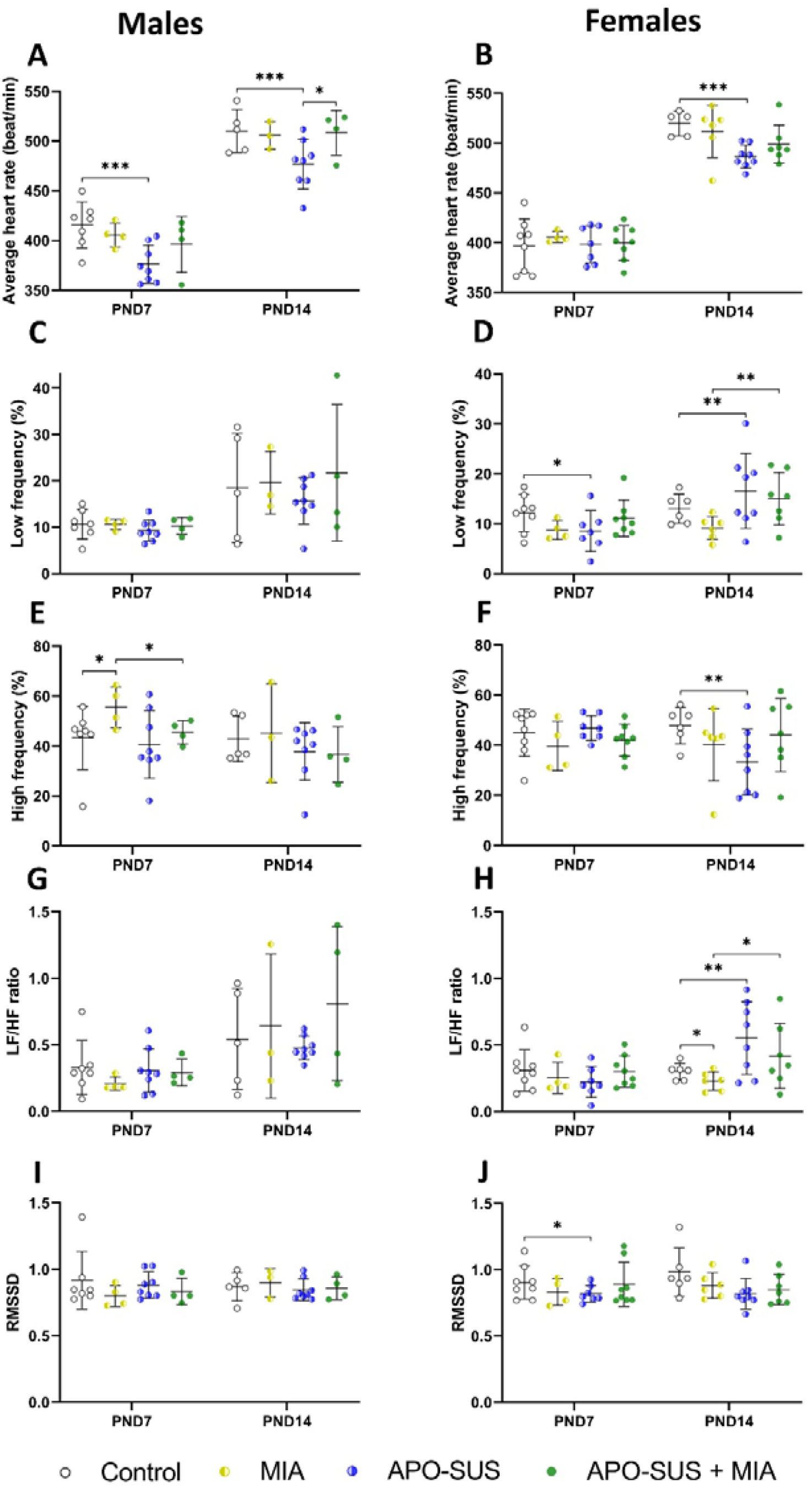
Heart rate variability in control, MIA, APO-SUS, and APO-SUS+MIA rats. Average heart rate (beat/min) in males (**A**.) and females (**B**.). Low frequency (%) in males (**C**.) and females (**D**.). High frequency (%) in males (**E**.) and females (**F**.). LF/HF ratio in males (**G**.) and females (**H**.). Roots mean square of successive differences (RMSSD) between normal heartbeats in males (**I**.) and females (**J**.). N=3-8 males and 4-8 females per group. Statistically significant differences between groups are indicated by asterisks: *p<0.05, **p<0.01, ***p<0.001. Data is presented as mean± SD. Significant differences between time points and between MIA and APO-SUS are not shown.

In males, GEE analysis revealed a main effect of Genotype on heart rate (p < 0.01) and HF (p < 0.028), and a Genotype × MIA interaction for heart rate (p = 0.019). Post-hoc tests showed that male APO-SUS rats had a lower heart rate than controls on PND7 and PND14 (p < 0.001), and lower than APO-SUS+MIA rats on PND14 (p < 0.05). On PND7, male MIA rats had higher HF than both control and APO-SUS+MIA rats (p < 0.05).

In females, GEE analysis revealed a main effect of Genotype on heart rate, LF/HF ratio, and RMSSD (p < 0.05). A Genotype × MIA interaction was found for HF (p = 0.004) and RMSSD (p = 0.013). Post-hoc comparisons showed that APO-SUS rats had a lower heart rate than controls on PND14 (p < 0.001). LF was lower in APO-SUS rats than controls on PND7, but higher on PND14 (p < 0.05). LF was also higher in APO-SUS+MIA rats than MIA rats on PND14 (p < 0.01). HF was lower in APO-SUS rats than controls on PND14 (p < 0.01). The LF/HF ratio was higher in APO-SUS rats than controls (p < 0.01), and lower in MIA rats compared to both control and APO-SUS+MIA rats (p < 0.05). RMSSD was reduced in APO-SUS rats compared to controls on PND7 (p < 0.05).

### 3.3. Adolescent control and APO-SUS rats exposed to a prenatal immune activation did not show anticipatory behavior

To determine baseline differences, we first measured locomotor activity and number of rearing between groups for the pre-anticipatory phase (days 5-6, Supplementary Figure 1). During adolescence, GEE analysis showed main effects of Genotype on distance traveled and rearing in both males and females (p < 0.05), and of MIA on rearing in both sexes (p < 0.05). In adulthood, Genotype had a main effect on distance traveled in males (p = 0.007) and on rearing in both males and females (p < 0.05). A main effect of MIA was also observed on rearing in adult females (p = 0.020). Adolescent female MIA, APO-SUS, and APO-SUS+MIA rats traveled more than controls (p < 0.001). In adolescent and adult males, APO-SUS+MIA rats traveled more than control and MIA rats (p < 0.05). MIA females had more rearing than controls and APO-SUS+MIA rats (p < 0.01), while APO-SUS males had fewer rearing than controls and APO-SUS+MIA rats (p < 0.05). Adult APO-SUS and APO-SUS+MIA males reared more than controls (p < 0.05).

To assess anticipatory pleasure (Figure 3), we next compared the total distance traveled and number of rearings between the pre-anticipation phase (days 5-6), and the anticipation phase (days 9-10) within groups.

**Figure 3.**
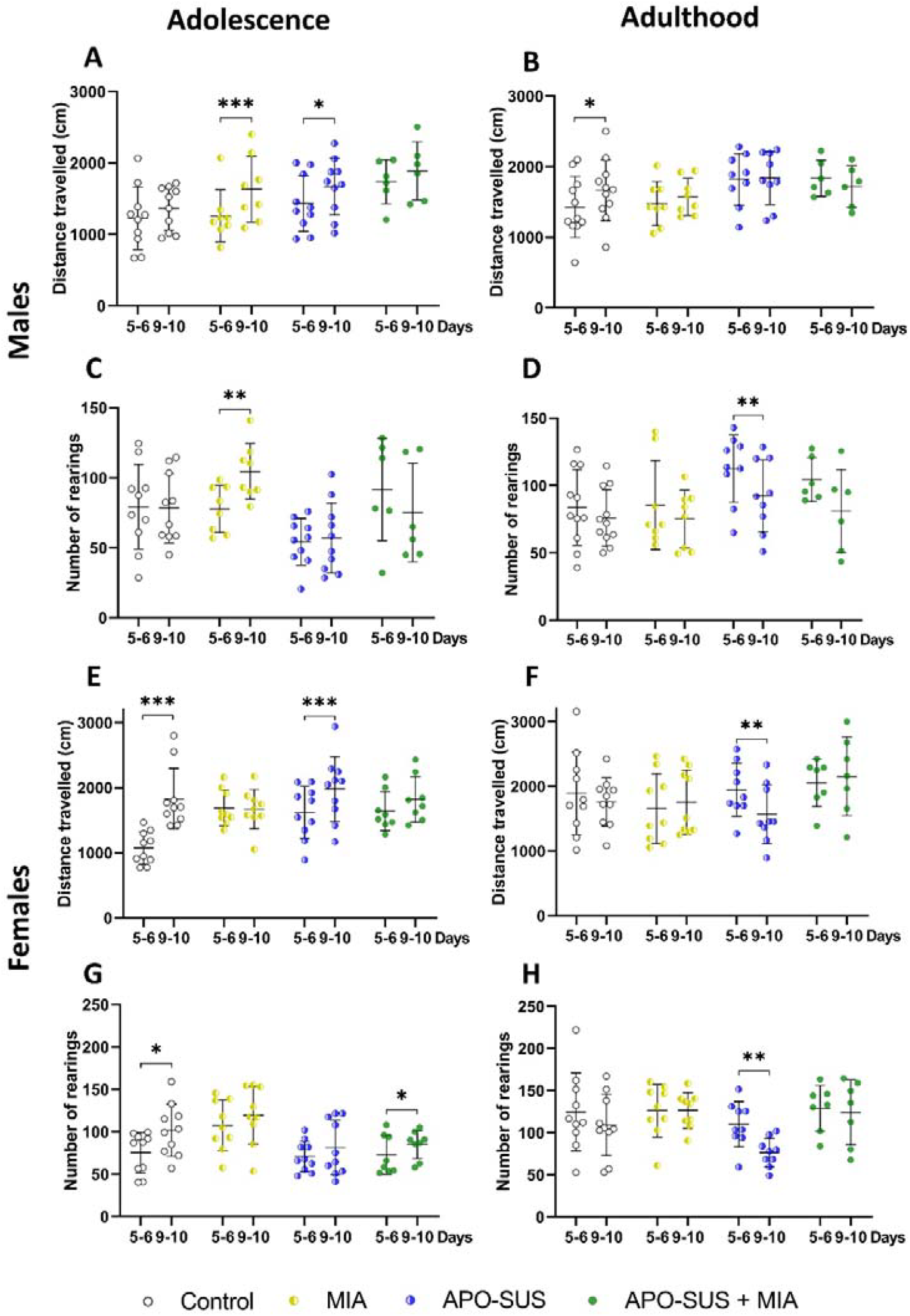
Within group comparison of the anticipatory locomotor and rearing activity in control, MIA, APO-SUS, and APO-SUS+MIA rats. Distance traveled in males during adolescence (**A**.) and adulthood (**B**.). Number of rearings in males during adolescence (**C**.) and adulthood (**D**.). Distance traveled in females during adolescence (**E**.) and adulthood (**F**.). Number of rearings in females during adolescence (**G**.) and adulthood (**H**.). Days 5–6 correspond to the pre-anticipation phase, and days 9– 10 to the anticipation phase. N=6-10 males and 7-10 females per group. Data is presented as mean±SD. Statistically significant differences within groups are indicated by asterisks: *p<0.05; **p<0.01, ***p<0.001. Significant differences between groups are not shown.

In adolescent males, GEE analysis revealed a main effect of Time for distance travelled, and Genotype × Time (p = 0.032) and Time × Genotype × MIA interactions (p = 0.011) for rearing. Post-hoc comparisons showed that MIA and APO-SUS rats traveled more on days 9–10 than days 5–6 (p < 0.05). MIA rats also showed increased rearing on days 9–10 (p = 0.006).

In adolescent females, GEE analysis showed significant main effect of Time (p < 0.001) and Time × MIA × Genotype interactions (p = 0.018) for distance travelled, and Time × MIA interactions (p < 0.001) for rearing. Control and APO-SUS+MIA rats showed increased distance travelled and rearing on days 9–10 compared to days 5-6 (p < 0.05).

In adult males, we observed a main effect of Time for rearing, and Genotype × Time interactions for distance traveled (p = 0.029). Control rats showed increased locomotor activity (p = 0.022), and APO-SUS rats showed a reduction in rearing (p < 0.001), on days 9–10 compared to days 5-6.

In adult females, a main effect of Time for rearing (p = 0.002), and Time × MIA interactions were observed for rearing and distance traveled (p < 0.001). APO-SUS rats showed reduced locomotor activity and rearing on days 9–10 compared to days 5–6 (p < 0.001).

### 3.4. APO-SUS and MIA rats showed elevated and reduced synaptophysin levels, respectively, and combined exposure normalized levels to controls

To assess synaptic alterations across development, synaptophysin and PSD95 protein levels were quantified in the hippocampus and frontal cortex during adolescence and adulthood (Figure 4).

**Figure 4.**
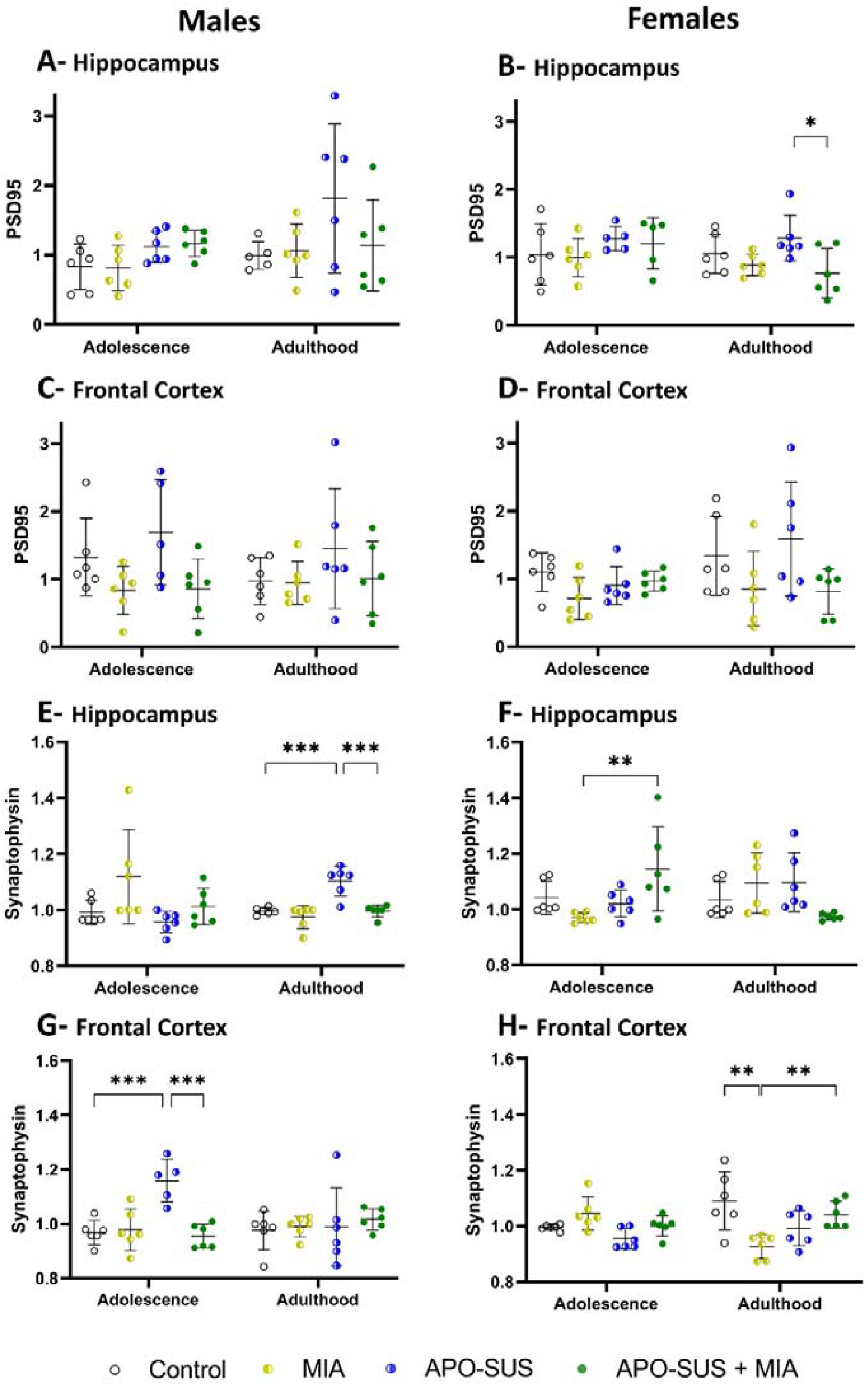
PSD95 and synaptophysin protein levels in control, MIA, APO-SUS, and APO-SUS+MIA rats. PSD95 levels in the hippocampus of males (**A**.) and females (**B**.). PSD95 levels in the frontal cortex of males (**C**.) and females (**D**.). Synaptophysin levels in the hippocampus of males (**E**.) and females (**F**.). Synaptophysin levels in the frontal cortex of males (**G**.) and females (**H**.).N=5-6 males and 5-6 females per group. Data is presented as mean±SD. Statistically significant differences within groups are indicated by asterisks: *p<0.05; **p<0.01, ***p<0.001. Significant differences between time points and between MIA and APO-SUS are not shown.

**Figure 5.**
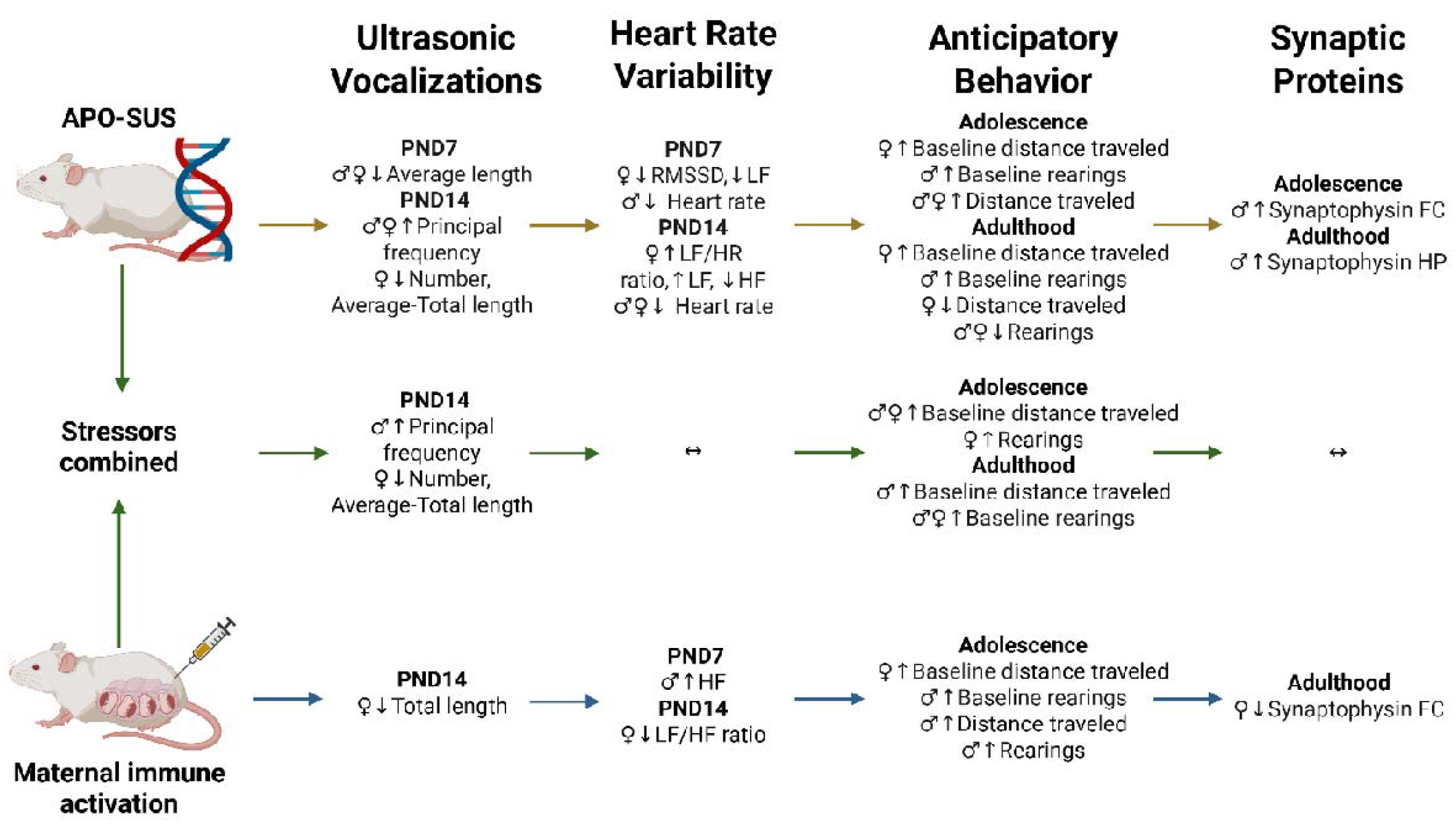
Summary figure of the effect of APO-SUS, MIA, and their combination on USVs, heart rate and behavior. Arrows up and down represent significant differences compared to control animals. HP=hippocampus, FC=frontal cortex ↔ = no significant changes compared with controls, ↑ = increase, ↓ = decrease.

#### Hippocampus

In adolescent males, ANOVA revealed a main effect of MIA on synaptophysin (p = 0.028), but post-hoc analysis did not show any significant group differences.

In adult males, main effects of MIA (p = 0.0003), Genotype (p = 0.0003), and a Genotype × MIA interaction (p = 0.009) were found on synaptophysin. Post-hoc tests showed that APO-SUS rats had higher synaptophysin levels than both control and APO-SUS+MIA rats (p < 0.001).

In adolescent females, a main effect of genotype (p = 0.040), and a Genotype × MIA interaction were observed for synaptophysin (p = 0.010). APO-SUS+MIA rats showed higher synaptophysin levels than MIA rats (p < 0.01).

In adult females, a Genotype × MIA interaction was found for synaptophysin (p = 0.014)), but post-hoc analysis did not show any significant group differences.

For PSD95, in adolescent males, a main effect of Genotype was found (p = 0.010),), but post-hoc analysis did not show any significant group differences.

In adult females, a main effect of MIA was observed for PSD95 (p = 0.011). Post-hoc analysis revealed that APO-SUS+MIA rats had lower PSD95 levels than APO-SUS rats (p = 0.03).

#### Frontal Cortex

In adolescent males, ANOVA revealed a main effect of MIA (p = 0.0014), Genotype (p = 0.0045), and a Genotype × MIA interaction (p = 0.0006) for synaptophysin levels. APO-SUS rats had higher levels than both control and APO-SUS+MIA rats (p < 0.001).

In adult males, no significant effects were detected.

In adolescent females, main effects of MIA (p = 0.008) and Genotype (p = 0.016) were found for synaptophysin, but post-hoc analysis did not show any significant group differences.

In adult females, a Genotype × MIA interaction was observed (p = 0.0012). MIA rats had lower synaptophysin levels than both control (p = 0.003) and APO-SUS+MIA rats (p = 0.044).

For PSD95, in adolescent males, a main effect of MIA was found in the frontal cortex (p = 0.009).

In adult females, a main effect of MIA was observed (p = 0.019), and in adolescent females, a Genotype × MIA interaction was found (p = 0.048) for PSD95, but post-hoc analysis did not show any significant group differences.

## 4. Discussion

In this study, we have innovatively combined APO-SUS rats with prenatal infection, exploring a new dimension of the multifactorial etiology of schizophrenia. Our findings reveal that APO-SUS rats exhibit early alterations in neurodevelopmental markers such as USVs and HRV, alongside age- and sex-dependent changes in synaptic and behavioral patterns (Figure 4). MIA interacted with this polygenic predisposition by preventing HRV reductions and normalizing synaptic density in APO-SUS rats, while also amplifying behavioral disruptions. This underscores the complex interplay between genetic vulnerabilities and environmental influences and highlights the necessity of exploring their intricate interplay to better understand the complex dynamics of neurodevelopmental disorders.

Regarding social-emotional communication (Brudzynski, 2013; Shair, 2007; Wöhr and Schwarting, 2013), we observed a main effect of genotype on USV parameters. APO-SUS rats, particularly females, exhibited shorter USV durations, fewer calls, and higher principal frequencies compared to control. These results align with increased high frequency calls and shortened call lengths in 12-day old pups exposed to maternal separation or genetic mutations related to neurodevelopmental disorders (Kaidbey et al., 2019; Lai et al., 2014; Roy et al., 2012; Schmeisser et al., 2012; Wöhr et al., 2022). These changes are consistent with the polyvagal hypothesis (Kaidbey et al., 2019) which proposes that acoustic signals reflect the physiological state. Low-frequency calls are longer and linked to calmer states, while shorter, higher-frequency calls have been associated with faster respirations rates and greater stress and anxiety—common comorbidities of schizophrenia (Braga et al., 2013; Kiran and Chaudhury, 2016). This interpretation aligns with altered HPA axis function previously reported in APO-SUS rats (Rots et al., 1996). These USV changes may represent an early manifestation of atypical affective states that could serve as markers for future communication deficits, often seen in schizophrenia patients (Walker et al., 2010)(Esposito et al., 2017). However, further studies are needed to determine whether these changes persist into adulthood.

In examining the dynamic interplay between the sympathetic and parasympathetic autonomic nervous systems (Kidwell and Ellenbroek, 2018; Rajendra Acharya et al., 2006), we observed that APO-SUS rats had a significantly lower average heart rate than controls on PND7 and 14. By PND14, female APO-SUS rats had a significantly reduced HRV compared to controls characterized by increased sympathetic dominance (elevated LF/HF ratio and LF percentage, alongside reduced HF percentage and RMSSD). This dysregulated autonomic nervous system function is associated with heightened stress sensitivity and commonly seen in many neuropsychiatric disorders (Benjamin et al., 2021; Haigh et al., 2021; Jung et al., 2019; Licht et al., 2009; Stogios et al., 2021). Female-specific reductions in HRV are consistent with the higher prevalence of anxiety and mood disorders in women and highlight the importance of sex as a biological variable in studying neurodevelopmental disorders (Ho et al., 2020). Overall, these results may indicate early dysregulation in the autonomic nervous system that might result in cognitive and emotional changes later in life (Thayer and Lane, 2000).

Consistent with previous research, APO-SUS rats exhibited higher baseline locomotor activity, likely reflecting heightened dopamine reactivity in the mesolimbic system—a known feature of schizophrenia (Cools et al., 1990; de Bruin et al., 2001). During adolescence, female control and APO-SUS rats, as well as male APO-SUS rats, showed anticipatory behavior (increased locomotion and/or rearings between pre- and post-anticipatory days). However, in adulthood, this behavior persisted only in male control rats, suggesting that APO-SUS rats developed apathy-like behaviors, aligning with the typical age of schizophrenia onset (Tandon et al., 2013). Notably, increased anticipatory locomotion was absent in adolescent male and adult female control rats, warranting further validation. The observed reduced locomotion and rearings in adult APO-SUS rats likely stemmed from insufficient motivation provided by the reward (sugary cereals), leading to habituation or boredom rather than aversion. Future studies could test food deprivation or stronger stimuli, like social interaction rewards.

Regarding prenatal immune activation, we did not see effects of poly I:C on the USVs of APO-SUS rats, which is consistent with studies reporting no influence of MIA on acoustic parameters like duration, bandwidth and peak frequency in rat pups (Möhrle et al., 2023; Potasiewicz et al., 2020). However, some studies found that MIA reduced the number and duration of calls, effects disappearing by PND14 (Malkova et al., 2012; Shinnyi Chou, Sean Jones, 2015; Wilkin-Krug et al., 2022), while others observed the opposite, with poly I:C exposure increasing call numbers (Schwartzer et al., 2013). These discrepancies may arise from differences in strain, sex, species, time, dose, administration route, and pup age (Jouda et al., 2019; Meyer et al., 2008, 2006). We observed an interaction between genotype and MIA in heart rate and HRV parameters. On PND7, MIA increased the high frequency of control, but not APO-SUS males. In females, APO-SUS effects on reducing HRV (elevated sympathetic dominance) were absent on PND14 when these APO-SUS rats were exposed to MIA. This suggests that MIA may have compensated for some of the effects of APO-SUS, possibly because MIA had opposite effects as it increased HRV (increased high frequency and reduced LF/HF) in controls. While MIA had no significant impact on USVs and appeared to restore HRV changes in APO-SUS rats, it inhibited the anticipatory locomotor increase observed in adolescent female control and male APO-SUS rats. Additionally, anticipatory behavior (increased locomotion and rearings) in adolescent males exposed to MIA disappeared in adulthood. These finding are complementary to other studies reporting anhedonia-like behavior in rats prenatally exposed to poly I:C (Khan et al., 2014; Reisinger et al., 2016), and align with clinical studies reporting disruptions in anticipatory pleasure in psychiatric disorders, such as major depressive disorder and schizophrenia (Frost and Strauss, 2016; Gard et al., 2007; Hallford et al., 2020; Mote et al., 2014; Sherdell et al., 2012).

Altered PSD95 and synaptophysin levels suggest disrupted synaptic function in both APO-SUS and MIA-exposed rats. The elevation of synaptophysin in the hippocampus during adulthood but not adolescence, and in the frontal cortex during adolescence, normalizing by adulthood, in male APO-SUS rats suggests altered and delayed synaptic pruning, resembling patterns observed in autism, where pruning peaks later than in healthy controls (Beopoulos et al., 2022). Delayed prefrontal cortex development is linked to poor executive function and psychiatric disorders in humans (Xie et al., 2023), aligning with previous studies showing executive dysfunction in APO-SUS rats (Maas et al., 2020). In contrast, female MIA rats exhibited the expected reduced PSD95 and synaptophysin levels in the frontal cortex during adulthood, indicating sex-specific vulnerability to prenatal infection. This reduction aligns with studies showing decreased synaptophysin and altered synaptic structure in the frontal cortex (LPS model) but contrasts with increased synaptophysin observed after poly I:C exposure in male rats, likely due to differences in MIA methods (Cieślik et al., 2020; Guerrin et al., 2023). Reduced PSD95 hippocampus levels in adult APO-SUS+MIA female rats, compared to APO-SUS alone, suggests a compounded synaptic dysfunction driven by genetic susceptibility and environmental factors. In contrast, synaptophysin levels in APO-SUS males normalized to control levels, highlighting the complex interplay of genetic and environmental influences in disrupting neurodevelopment.

These synaptic alterations may stem from abnormal immune system activity, particularly involving microglia and the complement system (Germann et al., 2021; Mattei et al., 2017). In support, studies link MIA to increased spine density and reduced CX3CR1 mRNA, a key regulator of synaptic pruning (Andoh et al., 2019; Fernández de Cossío et al., 2017). Excessive or reduced microglial activity has been shown to disrupt synaptic pruning (Cardozo et al., 2019; Germann et al., 2021; Han et al., 2022; Paolicelli et al., 2011), resulting in increased synaptic density in MIA, maternal separation and social defeat models (Dayananda et al., 2023; Guerrin et al., 2023; Mattei et al., 2017). Normalization of synaptophysin in male APO-SUS rats following MIA may results from long-term increases in microglial density and pruning activity, as seen in another study combining prenatal infection and adolescent stress (Guerrin et al., 2023). These findings align with the pathophysiology of schizophrenia and autism, where altered synaptic densities are prominent (Cardozo et al., 2019; Ebrahimi-Fakhari and Sahin, 2015; Gomot et al., 2008; Paolicelli et al., 2011). The hyperconnectivity in APO-SUS rats mirrors autism-related synaptic abnormalities (Ebrahimi-Fakhari and Sahin, 2015; Gomot et al., 2008; Hutsler and Zhang, 2010; Yizhar et al., 2011), while synaptic reductions in MIA rats resemble schizophrenia-associated deficits (Corradini et al., 2009; Onwordi et al., 2020; Osimo et al., 2019). However, we did not measure immune system activity nor synaptic pruning and longitudinal changes in synaptic density, which future research should investigate.

The observed protective effects of MIA suggest that it might not uniformly exacerbate alterations induced by APO-SUS but could also offer resilience. This aligns with existing research suggesting that MIA can sometimes play a protective role (Guerrin et al., 2021). For example, a previous study noted that MIA can prevent social defeat-induced anxiety and mitigate reductions in social behavior (Guerrin et al., 2023). Another study noted a protective effect of MIA against social isolation, associated with elevated levels of hippocampal oxytocin—a hormone that can reduce anxiety and social avoidance, notably by modulating glucocorticoid responses (Engelmann et al., 2004; Goh et al., 2020; Yoon and Kim, 2020). This mechanism may also underlie the protective effects observed in our study concerning HRV, as oxytocin has been demonstrated to enhance HRV through increased parasympathetic activity (Kemp et al., 2012; Martins et al., 2020; Schoormans et al., 2020), and is known to influence synaptic activity and plasticity (Rajamani et al., 2018). The concept of adversity enhancing resilience (Seery et al., 2013) aligns with our findings, suggesting that MIA can, under certain conditions, confer protective benefits in APO-SUS rats.

Importantly, the timing of poly I:C administration may have contributed to this protective outcome. Gestational day 15 represents a distinct neurodevelopmental window in rats, involving cortical neurogenesis, synaptogenesis, and microglial colonization, and differs substantially from earlier (GD9) or later (GD17) periods (Knuesel et al., 2014; Meyer, 2014; Meyer et al., 2007). Several studies using MIA on GD15 in rats, combined with adolescent stressors such as cannabis or methamphetamine exposure, social isolation, or repeated mild stress, have reported either no additive effects or even protective outcomes. These include reduced isolation-induced memory deficits, prevention of hyperlocomotion, prepulse inhibition, anxiety, and social withdrawal, and attenuation of dopaminergic, serotonergic, GABAergic, glutamatergic, and immune alterations (Dalton et al., 2012; Goh et al., 2020; Guerrin et al., 2021; Hollins et al., 2016; Lecca et al., 2019; Swanepoel et al., 2018; Yee et al., 2011). In contrast, studies in mice administering poly I:C on GD9 or GD17, in combination with adolescent stressors, Nurr1 suppression, or DISC1 mutations, have reported synergistic impairments in PPI, latent inhibition, cognitive flexibility, social interaction, and increased oxidative stress, dopaminergic dysfunction, and microglial activation (Deslauriers et al., 2014; Giovanoli et al., 2016, 2014, 2013; Guerrin et al., 2021; Lipina et al., 2013; Vuillermot et al., 2012). These findings suggest that MIA-induced vulnerability—or resilience—may be highly dependent on the timing of immune activation, the species used, and the nature of subsequent challenges. Future research directly comparing gestational timepoints is warranted to determine whether such outcomes are primarily timing-dependent or driven by interaction effects between developmental insult and genetic or environmental context.

This model has demonstrated a capacity to mimic certain neurodevelopmental phenotypes associated with psychiatric disorders like schizophrenia and autism, such as alterations in USVs, HRV, and synaptic density. These are influenced by both genetic predispositions of APO-SUS rats and environmental factors introduced through MIA. However, our study does not extend to measuring behaviors during adolescence and adulthood that are typically assessed in schizophrenia research, such as prepulse inhibition, latent inhibition, cognition functions and mood symptoms. Before this model can be effectively used to develop and test new therapeutic strategies aimed at modulating both the symptoms and underlying neurobiological disruptions seen in schizophrenia, future research should include these behavioral assessments. This additional data would allow to determine further whether MIA can exacerbate or mitigate these behavioral alterations, thus helping to ascertain the model’s behavioral relevance and enhancing its translational potential for human conditions.

Our results indicate that polygenic susceptibility in APO-SUS rats induces early neurodevelopmental changes, including altered USVs and HRV. These changes are indicative of increased sympathetic dominance, a condition commonly associated with psychiatric disorders. Interestingly, while prenatal immune activation did not affect USVs, it effectively mitigated APO-SUS-induced HRV reductions, increased synaptic density, and suppressed anticipatory locomotor increases in adolescence. These findings highlight the complex interplay between genetic predispositions and environmental factors. Future studies of genetic and environmental factors should further explore how early-life changes in USVs and HRV relate to later behavioral outcomes, as well as examine the role of synaptic pruning in the progression of neurodevelopmental disorders. Understanding these mechanisms is crucial for elucidating the genetic and environmental interactions on psychiatric conditions.

## Supporting information

MIA checklist

## 5. CRediT authorship contribution statement

**Kate Witt**: Data Curation, Investigation, Writing – Original Draft, Writing – Review & Editing. **Meyrick Kidwell**: Data Curation, Investigation, Writing – Review & Editing. **Janine Doorduin**: Conceptualization, Methodology, Supervision, Writing – Review & Editing. **Erik. F.J. de Vries**: Conceptualization, Methodology, Supervision, Project Administration, Writing – Review & Editing. **Iris E. Sommer:** Validation, Writing – Review & Editing. **Darren J. Day**: Methodology, Writing – Review & Editing. **Bart A. Ellenbroek:** Conceptualization, Methodology, Validation, Project Administration, Supervision, Writing – Review & Editing. **Cyprien G.J. Guerrin**: Conceptualization, Data Curation, Formal Analysis, Investigation, Methodology, Visualization, Writing – Original Draft, Writing – Review & Editing. All authors revised the manuscript for intellectual content and provided final approval for submission. All authors agree to be accountable for all aspects of the work, ensuring that questions related to accuracy or integrity are appropriately investigated and resolved.

## 6. Conflict of Interest

The authors have nothing to disclose.

## 7. Funding

This research did not receive any specific grant from funding agencies in the public, commercial, or not-for-profit sectors.

## Notes

### Competing Interest Statement

The authors have declared no competing interest.

## References

Alnæs, D., Kaufmann, T., Van Der Meer, D., Córdova-Palomera, A., Rokicki, J., Moberget, T., Bettella, F., Agartz, I., Barch, D.M., Bertolino, A., Brandt, C.L., Cervenka, S., Djurovic, S., Doan, N.T., Eisenacher, S., Fatouros-Bergman, H., Flyckt, L., Di Giorgio, A., Haatveit, B., Jönsson, E.G., Kirsch, P., Lund, M.J., Meyer-Lindenberg, A., Pergola, G., Schwarz, E., Smeland, O.B., Quarto, T., Zink, M., Andreassen, O.A., Westlye, L.T., 2019. Brain Heterogeneity in Schizophrenia and Its Association with Polygenic Risk. JAMA Psychiatry 76, 739–748. 10.1001/jamapsychiatry.2019.0257

Andoh, M., Shibata, K., Okamoto, K., Onodera, J., Morishita, K., Miura, Y., Ikegaya, Y., Koyama, R., 2019. Exercise Reverses Behavioral and Synaptic Abnormalities after Maternal Inflammation. Cell Rep. 27, 2817-2825.e5. 10.1016/j.celrep.2019.05.015

Barch, D.M., Ceaser, A., 2012. Cognition in Schizophrenia: Core Psychological and Neural Mechanisms Centrality of Cognition in Schizophrenia NIH Public Access. Trends Cogn Sci 16. 10.1016/j.tics.2011.11.015

Benjamin, B.R., Valstad, M., Elvsåshagen, T., Jönsson, E.G., Moberget, T., Winterton, A., Haram, M., Høegh, M.C., Lagerberg, T. V, Steen, N.E., Larsen, L., Andreassen, O.A., Westlye, L.T., Quintana, D.S., 2021. Heart rate variability is associated with disease severity in psychosis spectrum disorders. Prog. Neuro-Psychopharmacology Biol. Psychiatry 111, 110108. 10.1016/j.pnpbp.2020.110108

Beopoulos, A., Géa, M., Fasano, A., Iris, F., 2022. Autism spectrum disorders pathogenesis: Toward a comprehensive model based on neuroanatomic and neurodevelopment considerations. Front. Neurosci. 16, 988735. 10.3389/fnins.2022.988735

Braga, R.J., Reynolds, G.P., Siris, S.G., 2013. Anxiety comorbidity in schizophrenia. Psychiatry Res. 210, 1–7. 10.1016/j.psychres.2013.07.030

Brown, A.S., Meyer, U., 2018. Maternal immune activation and neuropsychiatric illness: A translational research perspective, American Journal of Psychiatry. 10.1176/appi.ajp.2018.17121311

Brudzynski, S.M., 2013. Ethotransmission: communication of emotional states through ultrasonic vocalization in rats. Curr. Opin. Neurobiol. 23, 310–317. 10.1016/j.conb.2013.01.014

Cardozo, P.L., de Lima, I.B.Q., Maciel, E.M.A., Silva, N.C., Dobransky, T., Ribeiro, F.M., 2019. Synaptic Elimination in Neurological Disorders. Curr. Neuropharmacol. 17, 1071–1095. 10.2174/1570159X17666190603170511

Careaga, M., Murai, T., Bauman, M.D., 2017. Maternal Immune Activation and Autism Spectrum Disorder: From Rodents to Nonhuman and Human Primates. Biol. Psychiatry 81, 391–401. 10.1016/j.biopsych.2016.10.020

Cieślik, M., Gąssowska-Dobrowolska, M., Jęśko, H., Czapski, G.A., Wilkaniec, A., Zawadzka, A., Dominiak, A., Polowy, R., Filipkowski, R.K., Boguszewski, P.M., Gewartowska, M., Frontczak-Baniewicz, M., Sun, G.Y., Beversdorf, D.Q., Adamczyk, A., 2020. Maternal Immune Activation Induces Neuroinflammation and Cortical Synaptic Deficits in the Adolescent Rat Offspring. Int. J. Mol. Sci. 10.3390/ijms21114097

Coley, A.A., Gao, W.-J., 2018. PSD95: A synaptic protein implicated in schizophrenia or autism? Prog. Neuro-Psychopharmacology Biol. Psychiatry 82, 187–194. 10.1016/j.pnpbp.2017.11.016

Coolen, M.W., van Loo, K.M.J., van Bakel, N.N.H.M., Ellenbroek, B.A., Cools, A.R., Martens, G.J.M., 2006. Reduced Aph-1b expression causes tissue- and substrate-specific changes in gamma-secretase activity in rats with a complex phenotype. FASEB J. Off. Publ. Fed. Am. Soc. Exp. Biol. 20, 175–177. 10.1096/fj.05-4337fje

Coolen, M.W., Van Loo, K.M.J., Van Bakel, N.N.H.M., Pulford, D.J., Serneels, L., De Strooper, B., Ellenbroek, B.A., Cools, A.R., Martens, G.J.M., 2005. Gene dosage effect on gamma-secretase component Aph-1b in a rat model for neurodevelopmental disorders. Neuron 45, 497–503. 10.1016/j.neuron.2004.12.054

Cools, A.R., Brachten, R., Heeren, D., Willemen, A., Ellenbroek, B., 1990. Search after neurobiological profile of individual-specific features of wistar rats. Brain Res. Bull. 24, 49–69. 10.1016/0361-9230(90)90288-B

Cools, A.R., Dierx, J., Coenders, C., Heeren, D., Ried, S., Jenks, B.G., Ellenbroek, B., 1993. Apomorphine-susceptible and apomorphine-unsusceptible Wistar rats differ in novelty-induced changes in hippocampal dynorphin B expression and two-way active avoidance: A new key in the search for the role of the hippocampal-accumbens axis. Behav. Brain Res. 55, 213–221. 10.1016/0166-4328(93)90117-9

Corradini, I., Verderio, C., Sala, M., Wilson, M.C., Matteoli, M., 2009. SNAP-25 in neuropsychiatric disorders. Ann. N. Y. Acad. Sci. 1152, 93–99. 10.1111/j.1749-6632.2008.03995.x

Dalton, V.S., Verdurand, M., Walker, A., Hodgson, D.M., Zavitsanou, K., 2012. Synergistic Effect between Maternal Infection and Adolescent Cannabinoid Exposure on Serotonin 5HT 1A Receptor Binding in the Hippocampus: Testing the “Two Hit” Hypothesis for the Development of Schizophrenia. ISRN Psychiatry 2012, 1–9. 10.5402/2012/451865

Dayananda, K.K., Ahmed, S., Wang, D., Polis, B., Islam, R., Kaffman, A., 2023. Early life stress impairs synaptic pruning in the developing hippocampus. Brain. Behav. Immun. 107, 16–31. 10.1016/j.bbi.2022.09.014

de Bruin, N.M., van Luijtelaar, E.L., Cools, A.R., Ellenbroek, B.A., 2001. Dopamine characteristics in rat genotypes with distinct susceptibility to epileptic activity: apomorphine-induced stereotyped gnawing and novelty/amphetamine-induced locomotor stimulation. Behav. Pharmacol. 12, 517–525. 10.1097/00008877-200111000-00013

Deslauriers, J., Racine, W., Sarret, P., Grignon, S., 2014. Preventive effect of α-lipoic acid on prepulse inhibition deficits in a juvenile two-hit model of schizophrenia. Neuroscience 272, 261–270. 10.1016/j.neuroscience.2014.04.061

Doorduin, J., de Vries, E.F.J., Willemsen, A.T.M., de Groot, J.C., Dierckx, R.A., Klein, H.C., 2009. Neuroinflammation in schizophrenia-related psychosis: a PET study. J. Nucl. Med. 50, 1801– 1807. 10.2967/jnumed.109.066647

Du, F., Cooper, A.J., Thida, T., Shinn, A.K., Cohen, B.M., Öngür, D., 2013. Myelin and axon abnormalities in schizophrenia measured with magnetic resonance imaging techniques. Biol. Psychiatry 74, 451–457. 10.1016/j.biopsych.2013.03.003

Ebrahimi-Fakhari, D., Sahin, M., 2015. Autism and the synapse: emerging mechanisms and mechanism-based therapies. Curr. Opin. Neurol. 28, 91–102. 10.1097/WCO.0000000000000186

Ellenbroek, B.A., Cools, A.R., 2002. Apomorphine susceptibility and animal models for psychopathology: Genes and environment. Behav. Genet. 10.1023/A:1020214322065

Ellenbroek, B.A., Cools, A.R., 2000. Animal models for the negative symptoms of schizophrenia. Behav. Pharmacol. 11, 223–233. 10.1097/00008877-200006000-00006

Ellenbroek, B.A., Geyer, M.A., Cools, A.R., 1995. The behavior of APO-SUS rats in animal models with construct validity for schizophrenia. J. Neurosci. 15, 7604–7611. 10.1523/jneurosci.15-11-07604.1995

Ellenbroek, B.A., Karl, T., 2016. Chapter 18 - Genetic Rat Models for Schizophrenia, in: Pletnikov, M. V, Waddington, J.L.B.T.-H. of B.N. (Eds.), Modeling the Psychopathological Dimensions of Schizophrenia. Elsevier, pp. 303–324. 10.1016/B978-0-12-800981-9.00018-3

Engelmann, M., Landgraf, R., Wotjak, C.T., 2004. The hypothalamic–neurohypophysial system regulates the hypothalamic–pituitary–adrenal axis under stress: An old concept revisited. Front. Neuroendocrinol. 25, 132–149. 10.1016/j.yfrne.2004.09.001

Esposito, G., Hiroi, N., Scattoni, M.L., 2017. Cry, Baby, Cry: Expression of Distress As a Biomarker and Modulator in Autism Spectrum Disorder. Int. J. Neuropsychopharmacol. 20, 498–503. 10.1093/ijnp/pyx014

Fernández de Cossío, L., Guzmán, A., van der Veldt, S., Luheshi, G.N., 2017. Prenatal infection leads to ASD-like behavior and altered synaptic pruning in the mouse offspring. Brain. Behav. Immun. 63, 88–98. 10.1016/j.bbi.2016.09.028

Fillman, S.G., Weickert, T.W., Lenroot, R.K., Catts, S. V., Bruggemann, J.M., Catts, V.S., Weickert, C.S., 2016. Elevated peripheral cytokines characterize a subgroup of people with schizophrenia displaying poor verbal fluency and reduced Broca’s area volume. Mol. Psychiatry 21, 1090– 1098. 10.1038/mp.2015.90

Flynn, S.W., Lang, D.J., Mackay, A.L., Goghari, V., Vavasour, I.M., Whittall, K.P., Smith, G.N., Arango, V., Mann, J.J., Dwork, A.J., Falkai, P., Honer, W.G., 2003. Abnormalities of myelination in schizophrenia detected in vivo with MRI, and post-mortem with analysis of oligodendrocyte proteins. Mol. Psychiatry 8, 811–820. 10.1038/sj.mp.4001337

Frost, K.H., Strauss, G.P., 2016. A Review of Anticipatory Pleasure in Schizophrenia. Curr. Behav. Neurosci. reports 3, 232–247. 10.1007/s40473-016-0082-5

Gard, D.E., Kring, A.M., Gard, M.G., Horan, W.P., Green, M.F., 2007. Anhedonia in schizophrenia: distinctions between anticipatory and consummatory pleasure. Schizophr. Res. 93, 253–260. 10.1016/j.schres.2007.03.008

Gasse, C., Wimberley, T., Wang, Y., Mors, O., Børglum, A., Damm Als, T., Werge, T., Nordentoft, M., Hougaard, D.M., Horsdal, H.T., 2019. Schizophrenia polygenic risk scores, urbanicity and treatment-resistant schizophrenia. 10.1016/j.schres.2019.08.008

Germann, M., Brederoo, S.G., Sommer, I.E.C., 2021. Abnormal synaptic pruning during adolescence underlying the development of psychotic disorders. Curr. Opin. Psychiatry 34, 222–227. 10.1097/YCO.0000000000000696

Giovanoli, S., Engler, H., Engler, A., Richetto, J., Feldon, J., Riva, M.A., Schedlowski, M., Meyer, U., 2016. Preventive effects of minocycline in a neurodevelopmental two-hit model with relevance to schizophrenia. Transl. Psychiatry 6, e772–9. 10.1038/tp.2016.38

Giovanoli, S., Engler, H., Engler, A., Richetto, J., Voget, M., Willi, R., Winter, C., Riva, M.A., Mortensen, P.B., Schedlowski, M., Meyer, U., 2013. Stress in puberty unmasks latent neuropathological consequences of prenatal immune activation in mice. Science (80-.). 339, 1100–1102. 10.1126/science.1228261

Giovanoli, S., Weber, L., Meyer, U., 2014. Single and combined effects of prenatal immune activation and peripubertal stress on parvalbumin and reelin expression in the hippocampal formation. Brain. Behav. Immun. 40, 48–54. 10.1016/j.bbi.2014.04.005

Glausier, J.R., Lewis, D.A., 2017. GABA and schizophrenia: Where we stand and where we need to go. Schizophr. Res. 181, 2–3. 10.1016/j.schres.2017.01.050

Goh, J.Y., O’Sullivan, S.E., Shortall, S.E., Zordan, N., Piccinini, A.M., Potter, H.G., Fone, K.C.F., King, M. V., 2020. Gestational poly(I:C) attenuates, not exacerbates, the behavioral, cytokine and mTOR changes caused by isolation rearing in a rat ‘dual-hit’ model for neurodevelopmental disorders. Brain. Behav. Immun. 89, 100–117. 10.1016/j.bbi.2020.05.076

Gomot, M., Belmonte, M.K., Bullmore, E.T., Bernard, F.A., Baron-Cohen, S., 2008. Brain hyper-reactivity to auditory novel targets in children with high-functioning autism. Brain 131, 2479– 2488. 10.1093/brain/awn172

Guerrin, C.G.J., Doorduin, J., Sommer, I.E., de Vries, E.F.J., 2021. The dual hit hypothesis of schizophrenia: Evidence from animal models. Neurosci. Biobehav. Rev. 131, 1150–1168. 10.1016/j.neubiorev.2021.10.025

Guerrin, C.G.J., Prasad, K., Vazquez-Matias, D.A., Zheng, J., Franquesa-Mullerat, M., Barazzuol, L., Doorduin, J., de Vries, E.F.J., 2023. Prenatal infection and adolescent social adversity affect microglia, synaptic density, and behavior in male rats. Neurobiol. Stress 27, 100580. 10.1016/j.ynstr.2023.100580

Guerrin, C.G.J., Shoji, A., Doorduin, J., de Vries, E.F.J., 2022. Immune Activation in Pregnant Rats Affects Brain Glucose Consumption, Anxiety-like Behaviour and Recognition Memory in their Male Offspring. Mol. Imaging Biol. 10.1007/s11307-022-01723-3

Haigh, S.M., Walford, T.P., Brosseau, P., 2021. Heart Rate Variability in Schizophrenia and Autism Front. Psychiatry

Hakak, Y., Walker, J.R., Li, C., Wong, W.H., Davis, K.L., Buxbaum, J.D., Haroutunian, V., Fienberg, A.A., 2001. Genome-wide expression analysis reveals dysregulation of myelination-related genes in chronic schizophrenia. Proc. Natl. Acad. Sci. U. S. A. 98, 4746–4751. 10.1073/pnas.081071198

Hallford, D.J., Barry, T.J., Austin, D.W., Raes, F., Takano, K., Klein, B., 2020. Impairments in episodic future thinking for positive events and anticipatory pleasure in major depression. J. Affect. Disord. 260, 536–543. 10.1016/j.jad.2019.09.039

Han, Q.-Q., Shen, S.-Y., Chen, X.-R., Pilot, A., Liang, L.-F., Zhang, J.-R., Li, W.-H., Fu, Y., Le, J.-M., Chen, P.-Q., Yu, J., 2022. Minocycline alleviates abnormal microglial phagocytosis of synapses in a mouse model of depression. Neuropharmacology 220, 109249. 10.1016/j.neuropharm.2022.109249

Hardie, I., Murray, A., King, J., Hall, H.A., Luedecke, E., Marryat, L., Thompson, L., Minnis, H., Wilson, P., Auyeung, B., 2025. Prenatal maternal infections and early childhood developmental outcomes: analysis of linked administrative health data for Greater Glasgow & Clyde, Scotland. J. Child Psychol. Psychiatry 66, 30–40. 10.1111/jcpp.14028

Ho, T.C., Pham, H.T., Miller, J.G., Kircanski, K., Gotlib, I.H., 2020. Sympathetic nervous system dominance during stress recovery mediates associations between stress sensitivity and social anxiety symptoms in female adolescents. Dev. Psychopathol. 32, 1914–1925. 10.1017/S0954579420001261

Hollins, S.L., Zavitsanou, K., Walker, F.R., Cairns, M.J., 2016. Alteration of transcriptional networks in the entorhinal cortex after maternal immune activation and adolescent cannabinoid exposure. Brain. Behav. Immun. 10.1016/j.bbi.2016.02.021

Howes, O.D., Murray, R.M., 2014. Schizophrenia: An integrated sociodevelopmental-cognitive model. Lancet. 10.1016/S0140-6736(13)62036-X

Hutsler, J.J., Zhang, H., 2010. Increased dendritic spine densities on cortical projection neurons in autism spectrum disorders. Brain Res. 1309, 83–94. 10.1016/j.brainres.2009.09.120

Jouda, J., Wöhr, M., Del Rey, A., 2019. Immunity and ultrasonic vocalization in rodents. Ann. N. Y. Acad. Sci. 1437, 68–82. 10.1111/nyas.13931

Jung, W., Jang, K.-I., Lee, S.-H., 2019. Heart and Brain Interaction of Psychiatric Illness: A Review Focused on Heart Rate Variability, Cognitive Function, and Quantitative Electroencephalography. Clin. Psychopharmacol. Neurosci. Off. Sci. J. Korean Coll. Neuropsychopharmacol. 17, 459–474. 10.9758/cpn.2019.17.4.459

Kaidbey, J.H., Ranger, M., Myers, M.M., Anwar, M., Ludwig, R.J., Schulz, A.M., Barone, J.L., Kolacz, J., Welch, M.G., 2019. Early Life Maternal Separation and Maternal Behaviour Modulate Acoustic Characteristics of Rat Pup Ultrasonic Vocalizations. Sci. Rep. 9, 19012. 10.1038/s41598-019-54800-z

Kemp, A.H., Quintana, D.S., Kuhnert, R.-L., Griffiths, K., Hickie, I.B., Guastella, A.J., 2012. Oxytocin Increases Heart Rate Variability in Humans at Rest: Implications for Social Approach-Related Motivation and Capacity for Social Engagement. PLoS One 7, e44014.

Khan, D., Fernando, P., Cicvaric, A., Berger, A., Pollak, A., Monje, F.J., Pollak, D.D., 2014. Long-term effects of maternal immune activation on depression-like behavior in the mouse. Transl. Psychiatry 4, e363–e363. 10.1038/tp.2013.132

Kidwell, M., Ellenbroek, B.A., 2018. Heart and soul: heart rate variability and major depression. Behav. Pharmacol. 29, 152–164. 10.1097/FBP.0000000000000387

Kiran, C., Chaudhury, S., 2016. Prevalence of comorbid anxiety disorders in schizophrenia. Ind. Psychiatry J. 25, 35–40. 10.4103/0972-6748.196045

Knuesel, I., Chicha, L., Britschgi, M., Schobel, S.A., Bodmer, M., Hellings, J.A., Toovey, S., Prinssen, E.P., 2014. Maternal immune activation and abnormal brain development across CNS disorders. Nat. Rev. Neurol. 10, 643–660. 10.1038/nrneurol.2014.187

Lai, J.K.Y., Sobala-Drozdowski, M., Zhou, L., Doering, L.C., Faure, P.A., Foster, J.A., 2014. Temporal and spectral differences in the ultrasonic vocalizations of fragile X knock out mice during postnatal development. Behav. Brain Res. 259, 119–130. 10.1016/j.bbr.2013.10.049

Lecca, S., Luchicchi, A., Scherma, M., Fadda, P., Muntoni, A.L., Pistis, M., 2019. Δ9-Tetrahydrocannabinol During Adolescence Attenuates Disruption of Dopamine Function Induced in Rats by Maternal Immune Activation. Front. Behav. Neurosci. 13, 1–8. 10.3389/fnbeh.2019.00202

Licht, C.M.M., de Geus, E.J.C., van Dyck, R., Penninx, B.W.J.H., 2009. Association between anxiety disorders and heart rate variability in The Netherlands Study of Depression and Anxiety (NESDA). Psychosom. Med. 71, 508–518. 10.1097/PSY.0b013e3181a292a6

Lipina, T. V., Zai, C., Hlousek, D., Roder, J.C., Wong, A.H.C., 2013. Maternal immune activation during gestation interacts with Disc1 point mutation to exacerbate schizophrenia-related behaviors in mice. J. Neurosci. 33, 7654–7666. 10.1523/JNEUROSCI.0091-13.2013

Maas, D.A., Eijsink, V.D., Spoelder, M., van Hulten, J.A., De Weerd, P., Homberg, J.R., Vallès, A., Nait-Oumesmar, B., Martens, G.J.M., 2020. Interneuron hypomyelination is associated with cognitive inflexibility in a rat model of schizophrenia. Nat. Commun. 11, 1–16. 10.1038/s41467-020-16218-4

Malkova, N. V., Yu, C.Z., Hsiao, E.Y., Moore, M.J., Patterson, P.H., 2012. Maternal immune activation yields offspring displaying mouse versions of the three core symptoms of autism. Brain. Behav. Immun. 26, 607–616. 10.1016/j.bbi.2012.01.011

Martins, D., Davies, C., De Micheli, A., Oliver, D., Krawczun-Rygmaczewska, A., Fusar-Poli, P., Paloyelis, Y., 2020. Intranasal oxytocin increases heart-rate variability in men at clinical high risk for psychosis: a proof-of-concept study. Transl. Psychiatry 10, 227. 10.1038/s41398-020-00890-7

Mattei, D., Ivanov, A., Ferrai, C., Jordan, P., Guneykaya, D., Buonfiglioli, A., Schaafsma, W., Przanowski, P., Deuther-Conrad, W., Brust, P., Hesse, S., Patt, M., Sabri, O., Ross, T.L., Eggen, B.J.L., Boddeke, E.W.G.M., Kaminska, B., Beule, D., Pombo, A., Kettenmann, H., Wolf, S.A., 2017. Maternal immune activation results in complex microglial transcriptome signature in the adult offspring that is reversed by minocycline treatment. Transl. Psychiatry 7. 10.1038/tp.2017.80

Meyer, U., 2014. Prenatal Poly(I:C) exposure and other developmental immune activation models in rodent systems. Biol. Psychiatry 75, 307–315. 10.1016/j.biopsych.2013.07.011

Meyer, U., Murray, P.J., Urwyler, A., Yee, B.K., Schedlowski, M., Feldon, J., 2008. Adult behavioral and pharmacological dysfunctions following disruption of the fetal brain balance between pro-inflammatory and IL-10-mediated anti-inflammatory signaling. Mol. Psychiatry 13, 208–221. 10.1038/sj.mp.4002042

Meyer, U., Nyffeler, M., Engler, A., Urwyler, A., Schedlowski, M., Knuesel, I., Yee, B.K., Feldon, J., 2006. The time of prenatal immune challenge determines the specificity of inflammation-mediated brain and behavioral pathology. J. Neurosci. 26, 4752–4762. 10.1523/JNEUROSCI.0099-06.2006

Meyer, U., Yee, B.K., Feldon, J., 2007. The Neurodevelopmental Impact of Prenatal Infections at Different Times of Pregnancy: The Earlier the Worse? Neurosci. 13, 241–256. 10.1177/1073858406296401

Modai, S., Shomron, N., 2016. Molecular Risk Factors for Schizophrenia. Trends Mol. Med. 22, 242– 253. 10.1016/j.molmed.2016.01.006

Möhrle, D., Yuen, M., Zheng, A., Haddad, F.L., Allman, B.L., Schmid, S., 2023. Characterizing maternal isolation-induced ultrasonic vocalizations in a gene–environment interaction rat model for autism. Genes, Brain Behav. n/a, e12841. 10.1111/gbb.12841

Mote, J., Minzenberg, M.J., Carter, C.S., Kring, A.M., 2014. Deficits in anticipatory but not consummatory pleasure in people with recent-onset schizophrenia spectrum disorders. Schizophr. Res. 159, 76–79. 10.1016/j.schres.2014.07.048

Murray, K.N., Edye, M.E., Manca, M., Vernon, A.C., Oladipo, J.M., Fasolino, V., Harte, M.K., Mason, V., Grayson, B., McHugh, P.C., Knuesel, I., Prinssen, E.P., Hager, R., Neill, J.C., 2019. Evolution of a maternal immune activation (mIA) model in rats: Early developmental effects. Brain. Behav. Immun. 75, 48–59. 10.1016/j.bbi.2018.09.005

Ng, F., Berk, M., Dean, O., Bush, A.I., 2008. Oxidative stress in psychiatric disorders: Evidence base and therapeutic implications. Int. J. Neuropsychopharmacol. 10.1017/S1461145707008401

Oliveras, I., Cañete, T., Sampedro-Viana, D., Río-Álamos, C., Tobeña, A., Corda, M.G., Giorgi, O., Fernández-Teruel, A., 2023. Neurobehavioral Profiles of Six Genetically-based Rat Models of Schizophrenia-related Symptoms. Curr. Neuropharmacol. 21, 1934–1952. 10.2174/1570159X21666230221093644

Onwordi, E.C., Halff, E.F., Whitehurst, T., Mansur, A., Cotel, M.-C., Wells, L., Creeney, H., Bonsall, D., Rogdaki, M., Shatalina, E., Reis Marques, T., Rabiner, E.A., Gunn, R.N., Natesan, S., Vernon, A.C., Howes, O.D., 2020. Synaptic density marker SV2A is reduced in schizophrenia patients and unaffected by antipsychotics in rats. Nat. Commun. 11, 246. 10.1038/s41467-019-14122-0

Osimo, E.F., Beck, K., Reis Marques, T., Howes, O.D., 2019. Synaptic loss in schizophrenia: a meta-analysis and systematic review of synaptic protein and mRNA measures. Mol. Psychiatry 24, 549–561. 10.1038/s41380-018-0041-5

Paolicelli, R.C., Bolasco, G., Pagani, F., Maggi, L., Scianni, M., Panzanelli, P., Giustetto, M., Ferreira, T.A., Guiducci, E., Dumas, L., Ragozzino, D., Gross, C.T., 2011. Synaptic Pruning by Microglia Is Necessary for Normal Brain Development. Science (80-.). 333, 1456–1458. 10.1126/science.1202529

Potasiewicz, A., Gzielo, K., Popik, P., Nikiforuk, A., 2020. Effects of prenatal exposure to valproic acid or poly(I:C) on ultrasonic vocalizations in rat pups: The role of social cues. Physiol. Behav. 225, 113113. 10.1016/j.physbeh.2020.113113

Rajamani, K.T., Wagner, S., Grinevich, V., Harony-Nicolas, H., 2018. Oxytocin as a Modulator of Synaptic Plasticity: Implications for Neurodevelopmental Disorders. Front. Synaptic Neurosci. 10.

Rajasekaran, A., Venkatasubramanian, G., Berk, M., Debnath, M., 2015. Mitochondrial dysfunction in schizophrenia: pathways, mechanisms and implications. Neurosci. Biobehav. Rev. 48, 10–21. 10.1016/j.neubiorev.2014.11.005

Rajendra Acharya, U., Paul Joseph, K., Kannathal, N., Lim, C.M., Suri, J.S., 2006. Heart rate variability: a review. Med. Biol. Eng. Comput. 44, 1031–1051. 10.1007/s11517-006-0119-0

Reisinger, S.N., Kong, E., Khan, D., Schulz, S., Ronovsky, M., Berger, S., Horvath, O., Cabatic, M., Berger, A., Pollak, D.D., 2016. Maternal immune activation epigenetically regulates hippocampal serotonin transporter levels. Neurobiol. Stress 4, 34–43. 10.1016/j.ynstr.2016.02.007

Riley, B., Kendler, K.S., 2006. Molecular genetic studies of schizophrenia. Eur. J. Hum. Genet. 14, 669– 680. 10.1038/sj.ejhg.5201571

Ripke, S., Neale, B.M., Corvin, A., Walters, J.T.R., Farh, K.H., Holmans, P.A., Lee, P., Bulik-Sullivan, B., Collier, D.A., Huang, H., Pers, T.H., Agartz, I., Agerbo, E., Albus, M., Alexander, M., Amin, F., Bacanu, S.A., Begemann, M., Belliveau, R.A., Bene, J., Bergen, S.E., Bevilacqua, E., Bigdeli, T.B., Black, D.W., Bruggeman, R., Buccola, N.G., Buckner, R.L., Byerley, W., Cahn, W., Cai, G., Campion, D., Cantor, R.M., Carr, V.J., Carrera, N., Catts, S. V., Chambert, K.D., Chan, R.C.K., Chen, R.Y.L., Chen, E.Y.H., Cheng, W., Cheung, E.F.C., Chong, S.A., Cloninger, C.R., Cohen, D., Cohen, N., Cormican, P., Craddock, N., Crowley, J.J., Curtis, D., Davidson, M., Davis, K.L., Degenhardt, F., Del Favero, J., Demontis, D., Dikeos, D., Dinan, T., Djurovic, S., Donohoe, G., Drapeau, E., Duan, J., Dudbridge, F., Durmishi, N., Eichhammer, P., Eriksson, J., Escott-Price, V., Essioux, L., Fanous, A.H., Farrell, M.S., Frank, J., Franke, L., Freedman, R., Freimer, N.B., Friedl, M., Friedman, J.I., Fromer, M., Genovese, G., Georgieva, L., Giegling, I., Giusti-Rodríguez, P., Godard, S., Goldstein, J.I., Golimbet, V., Gopal, S., Gratten, J., De Haan, L., Hammer, C., Hamshere, M.L., Hansen, M., Hansen, T., Haroutunian, V., Hartmann, A.M., Henskens, F.A., Herms, S., Hirschhorn, J.N., Hoffmann, P., Hofman, A., Hollegaard, M. V., Hougaard, D.M., Ikeda, M., Joa, I., Julià, A., Kahn, R.S., Kalaydjieva, L., Karachanak-Yankova, S., Karjalainen, J., Kavanagh, D., Keller, M.C., Kennedy, J.L., Khrunin, A., Kim, Y., Klovins, J., Knowles, J.A., Konte, B., Kucinskas, V., Kucinskiene, Z.A., Kuzelova-Ptackova, H., Kähler, A.K., Laurent, C., Keong, J.L.C., Lee, S.H., Legge, S.E., Lerer, B., Li, M., Li, T., Liang, K.Y., Lieberman, J., Limborska, S., Loughland, C.M., Lubinski, J., Lönnqvist, J., Macek, M., Magnusson, P.K.E., Maher, B.S., Maier, W., Mallet, J., Marsal, S., Mattheisen, M., Mattingsdal, M., McCarley, R.W., McDonald, C., McIntosh, A.M., Meier, S., Meijer, C.J., Melegh, B., Melle, I., Mesholam-Gately, R.I., Metspalu, A., Michie, P.T., Milani, L., Milanova, V., Mokrab, Y., Morris, D.W., Mors, O., Murphy, K.C., Murray, R.M., Myin-Germeys, I., Müller-Myhsok, B., Nelis, M., Nenadic, I., Nertney, D.A., Nestadt, G., Nicodemus, K.K., Nikitina-Zake, L., Nisenbaum, L., Nordin, A., O’Callaghan, E., O’Dushlaine, C., O’Neill, F.A., Oh, S.Y., Olincy, A., Olsen, L., Van Os, J., Pantelis, C., Papadimitriou, G.N., Papiol, S., Parkhomenko, E., Pato, M.T., Paunio, T., Pejovic-Milovancevic, M., Perkins, D.O., Pietiläinen, O., Pimm, J., Pocklington, A.J., Powell, J., Price, A., Pulver, A.E., Purcell, S.M., Quested, D., Rasmussen, H.B., Reichenberg, A., Reimers, M.A., Richards, A.L., Roffman, J.L., Roussos, P., Ruderfer, D.M., Salomaa, V., Sanders, A.R., Schall, U., Schubert, C.R., Schulze, T.G., Schwab, S.G., Scolnick, E.M., Scott, R.J., Seidman, L.J., Shi, J., Sigurdsson, E., Silagadze, T., Silverman, J.M., Sim, K., Slominsky, P., Smoller, J.W., So, H.C., Spencer, C.C.A., Stahl, E.A., Stefansson, H., Steinberg, S., Stogmann, E., Straub, R.E., Strengman, E., Strohmaier, J., Stroup, T.S., Subramaniam, M., Suvisaari, J., Svrakic, D.M., Szatkiewicz, J.P., Söderman, E., Thirumalai, S., Toncheva, D., Tosato, S., Veijola, J., Waddington, J., Walsh, D., Wang, D., Wang, Q., Webb, B.T., Weiser, M., Wildenauer, D.B., Williams, N.M., Williams, S., Witt, S.H., Wolen, A.R., Wong, E.H.M., Wormley, B.K., Xi, H.S., Zai, C.C., Zheng, X., Zimprich, F., Wray, N.R., Stefansson, K., Visscher, P.M., Adolfsson, R., Andreassen, O.A., Blackwood, D.H.R., Bramon, E., Buxbaum, J.D., Børglum, A.D., Cichon, S., Darvasi, A., Domenici, E., Ehrenreich, H., Esko, T., Gejman, P. V., Gill, M., Gurling, H., Hultman, C.M., Iwata, N., Jablensky, A. V., Jönsson, E.G., Kendler, K.S., Kirov, G., Knight, J., Lencz, T., Levinson, D.F., Li, Q.S., Liu, J., Malhotra, A.K., McCarroll, S.A., McQuillin, A., Moran, J.L., Mortensen, P.B., Mowry, B.J., Nöthen, M.M., Ophoff, R.A., Owen, M.J., Palotie, A., Pato, C.N., Petryshen, T.L., Posthuma, D., Rietschel, M., Riley, B.P., Rujescu, D., Sham, P.C., Sklar, P., St Clair, D., Weinberger, D.R., Wendland, J.R., Werge, T., Daly, M.J., Sullivan, P.F., O’Donovan, M.C., 2014. Biological insights from 108 schizophrenia-associated genetic loci. Nature 511, 421–427. 10.1038/nature13595

Rots, N.Y., Cools, A.R., Bérod, A., Voorn, P., Rostène, W., De Kloet, E.R., 1996. Rats bred for enhanced apomorphine susceptibility have elevated tyrosine hydroxylase mRNA and dopamine D2-receptor binding sites in nigrostriatal and tuberoinfundibular dopamine systems. Brain Res. 710, 189–196. 10.1016/0006-8993(95)01379-2

Roy, S., Watkins, N., Heck, D., 2012. Comprehensive Analysis of Ultrasonic Vocalizations in a Mouse Model of Fragile X Syndrome Reveals Limited, Call Type Specific Deficits. PLoS One 7, e44816.

Sarkar, T., Patro, N., Patro, I.K., 2019. Cumulative multiple early life hits-a potent threat leading to neurological disorders. Brain Res. Bull. 147, 58–68. 10.1016/j.brainresbull.2019.02.005

Schmeisser, M.J., Ey, E., Wegener, S., Bockmann, J., Stempel, A.V., Kuebler, A., Janssen, A.-L., Udvardi, P.T., Shiban, E., Spilker, C., Balschun, D., Skryabin, B. V, Dieck, S.tom, Smalla, K.-H., Montag, D., Leblond, C.S., Faure, P., Torquet, N., Le Sourd, A.-M., Toro, R., Grabrucker, A.M., Shoichet, S.A., Schmitz, D., Kreutz, M.R., Bourgeron, T., Gundelfinger, E.D., Boeckers, T.M., 2012. Autistic-like behaviours and hyperactivity in mice lacking ProSAP1/Shank2. Nature 486, 256–260. 10.1038/nature11015

Schmidt, M.J., Mirnics, K., 2014. Neurodevelopment, GABA System Dysfunction, and Schizophrenia. Neuropsychopharmacology 40, 190–206. 10.1038/npp.2014.95

Schoormans, D., Kop, W.J., Kunst, L.E., Riem, M.M.E., 2020. Oxytocin effects on resting-state heart rate variability in women: The role of childhood rearing experiences. Compr. Psychoneuroendocrinology 3, 100007. 10.1016/j.cpnec.2020.100007

Schwartzer, J.J., Careaga, M., Onore, C.E., Rushakoff, J.A., Berman, R.F., Ashwood, P., 2013. Maternal immune activation and strain specific interactions in the development of autism-like behaviors in mice. Transl. Psychiatry 3, e240–e240. 10.1038/tp.2013.16

Seery, M.D., Leo, R.J., Lupien, S.P., Kondrak, C.L., Almonte, J.L., 2013. An Upside to Adversity?: Moderate Cumulative Lifetime Adversity Is Associated With Resilient Responses in the Face of Controlled Stressors. Psychol. Sci. 24, 1181–1189. 10.1177/0956797612469210

Sekar, A., Bialas, A.R., de Rivera, H., Davis, A., Hammond, T.R., Kamitaki, N., Tooley, K., Presumey, J., Baum, M., Van Doren, V., Genovese, G., Rose, S.A., Handsaker, R.E., Daly, M.J., Carroll, M.C., Stevens, B., McCarroll, S.A., Consortium, S.W.G. of the P.G., 2016. Schizophrenia risk from complex variation of complement component 4. Nature 530, 177–183. 10.1038/nature16549

Selten, M.M., Meyer, F., Ba, W., Vallès, A., Maas, D.A., Negwer, M., Eijsink, V.D., Van Vugt, R.W.M., Van Hulten, J.A., Van Bakel, N.H.M., Roosen, J., Van Der Linden, R.J., Schubert, D., Verheij, M.M.M., Kasri, N.N., Martens, G.J.M., 2016. Increased GABA B receptor signaling in a rat model for schizophrenia. Sci. Rep. 6. 10.1038/srep34240

Shair, H.N., 2007. Acquisition and expression of a socially mediated separation response. Behav. Brain Res. 182, 180–192. 10.1016/j.bbr.2007.02.016

Sherdell, L., Waugh, C.E., Gotlib, I.H., 2012. Anticipatory pleasure predicts motivation for reward in major depression. J. Abnorm. Psychol. 121, 51–60. 10.1037/a0024945

Shinnyi Chou, Sean Jones, and M.L., 2015. Adolescent olanzapine sensitization is correlated with hippocampal stem cell proliferation in a maternal immune activation rat model of schizophrenia. Brain Res. 176, 139–148. 10.1016/j.brainres.2015.05.036.Adolescent

Sommer, I.E., Tiihonen, J., van Mourik, A., Tanskanen, A., Taipale, H., 2020. The clinical course of schizophrenia in women and men—a nation-wide cohort study. npj Schizophr. 6. 10.1038/s41537-020-0102-z

Stogios, N., Gdanski, A., Gerretsen, P., Chintoh, A.F., Graff-Guerrero, A., Rajji, T.K., Remington, G., Hahn, M.K., Agarwal, S.M., 2021. Autonomic nervous system dysfunction in schizophrenia: impact on cognitive and metabolic health. NPJ Schizophr. 7, 22. 10.1038/s41537-021-00151-6

Swanepoel, T., Möller, M., Harvey, B.H., 2018. N-acetyl cysteine reverses bio-behavioural changes induced by prenatal inflammation, adolescent methamphetamine exposure and combined challenges. Psychopharmacology (Berl). 235, 351–368. 10.1007/s00213-017-4776-5

Tandon, R., Gaebel, W., Barch, D.M., Bustillo, J., Gur, R.E., Heckers, S., Malaspina, D., Owen, M.J., Schultz, S., Tsuang, M., Van Os, J., Carpenter, W., 2013. Definition and description of schizophrenia in the DSM-5. 10.1016/j.schres.2013.05.028

Thayer, J.F., Lane, R.D., 2000. A model of neurovisceral integration in emotion regulation and dysregulation. J. Affect. Disord. 61, 201–216. 10.1016/s0165-0327(00)00338-4

Uher, R., 2014. Gene-environment interactions in severe mental illness. Front. Psychiatry. 10.3389/fpsyt.2014.00048

van der Elst, M.C.J., Ellenbroek, B.A., Cools, A.R., 2006. Cocaine strongly reduces prepulse inhibition in apomorphine-susceptible rats, but not in apomorphine-unsusceptible rats: Regulation by dopamine D2 receptors. Behav. Brain Res. 175, 392–398. 10.1016/j.bbr.2006.09.014

van Loo, K.M.J., Martens, G.J.M., 2007. Identification of genetic and epigenetic variations in a rat model for neurodevelopmental disorders. Behav. Genet. 37, 697–705. 10.1007/s10519-007-9164-1

Van Schijndel, J.E., Van Zweeden, M., Van Loo, K.M.J., Martens, G.J.M., 2010. Gene expression profiling in brain regions of a rat model displaying schizophrenia-related features. Behav. Brain Res. 207, 476–479. 10.1016/j.bbr.2009.10.042

van Vugt, R.W.M., Meyer, F., van Hulten, J.A., Vernooij, J., Cools, A.R., Verheij, M.M.M., Martens, G.J.M., 2014. Maternal care affects the phenotype of a rat model for schizophrenia. Front. Behav. Neurosci. 8. 10.3389/fnbeh.2014.00268

Vuillermot, S., Joodmardi, E., Perlmann, T., Ögren, S.O., Feldon, J., Meyer, U., 2012. Prenatal immune activation interacts with Genetic Nurr1 deficiency in the development of attentional impairments. J. Neurosci. 32, 436–451. 10.1523/JNEUROSCI.4831-11.2012

Walker, E., Shapiro, D., Esterberg, M., Trotman, H., 2010. Neurodevelopment and Schizophrenia: Broadening the Focus. Curr. Dir. Psychol. Sci. 19, 204–208. 10.1177/0963721410377744

Wiedenmann, B., Franke, W.W., Kuhn, C., Moll, R., Gould, V.E., 1986. Synaptophysin: a marker protein for neuroendocrine cells and neoplasms. Proc. Natl. Acad. Sci. 83, 3500–3504. 10.1073/pnas.83.10.3500

Wilkin-Krug, L.C.M., Macaskill, A.C., Ellenbroek, B.A., 2022. Preweaning environmental enrichment alters neonatal ultrasonic vocalisations in a rat model for prenatal infections. Behav. Pharmacol. 33, 402–417. 10.1097/FBP.0000000000000688

Wöhr, M., Fong, W.M., Janas, J.A., Mall, M., Thome, C., Vangipuram, M., Meng, L., Südhof, T.C., Wernig, M., 2022. Myt1l haploinsufficiency leads to obesity and multifaceted behavioral alterations in mice. Mol. Autism 13, 19. 10.1186/s13229-022-00497-3

Wöhr, M., Schwarting, R.K.W., 2013. Affective communication in rodents: ultrasonic vocalizations as a tool for research on emotion and motivation. Cell Tissue Res. 354, 81–97. 10.1007/s00441-013-1607-9

Xie, C., Xiang, S., Shen, C., Peng, X., Kang, J., Li, Y., Cheng, W., He, S., Bobou, M., Broulidakis, M.J., van Noort, B.M., Zhang, Z., Robinson, L., Vaidya, N., Winterer, J., Zhang, Y., King, S., Banaschewski, T., Barker, G.J., Bokde, A.L.W., Bromberg, U., Büchel, C., Flor, H., Grigis, A., Garavan, H., Gowland, P., Heinz, A., Ittermann, B., Lemaître, H., Martinot, J.-L., Martinot, M.-L.P., Nees, F., Orfanos, D.P., Paus, T., Poustka, L., Fröhner, J.H., Schmidt, U., Sinclair, J., Smolka, M.N., Stringaris, A., Walter, H., Whelan, R., Desrivières, S., Sahakian, B.J., Robbins, T.W., Schumann, G., Jia, T., Feng, J., van Noort, B.M., Consortium, I., Consortium, S., Consortium, Z.I.B., 2023. A shared neural basis underlying psychiatric comorbidity. Nat. Med. 29, 1232–1242. 10.1038/s41591-023-02317-4

Yee, N., Ribic, A., de Roo, C.C., Fuchs, E., 2011. Differential effects of maternal immune activation and juvenile stress on anxiety-like behaviour and physiology in adult rats: No evidence for the “double-hit hypothesis.” Behav. Brain Res. 224, 180–188. 10.1016/j.bbr.2011.05.040

Yizhar, O., Fenno, L.E., Prigge, M., Schneider, F., Davidson, T.J., O’Shea, D.J., Sohal, V.S., Goshen, I., Finkelstein, J., Paz, J.T., Stehfest, K., Fudim, R., Ramakrishnan, C., Huguenard, J.R., Hegemann, P., Deisseroth, K., 2011. Neocortical excitation/inhibition balance in information processing and social dysfunction. Nature 477, 171–178. 10.1038/nature10360

Yoon, S., Kim, Y.-K., 2020. The Role of the Oxytocin System in Anxiety Disorders BT -Anxiety Disorders: Rethinking and Understanding Recent Discoveries, in: Kim, Y.-K. (Ed.),. Springer Singapore, Singapore, pp. 103–120. 10.1007/978-981-32-9705-0_7

