## Supplementary Materials for "Apomorphine susceptibility and prenatal infection alter neurodevelopment, synaptic density and anticipatory behavior in rats"

### 2.1. Animals

All experiments were conducted in accordance with Victoria University of Wellington’s animal care principles and were approved by the Victoria University of Wellington’s Animal Ethics Committee. Normal Wistar and APO-SUS rats were used. APO-SUS rats are Wistar rats that have been selectively bred based on their behavioral susceptibility to an injection of apomorphine. In particular, Wistar rats displaying stereotypic gnawing behavior (>500 gnaws in 45 min) upon subcutaneous injection of 1.5 mg/kg apomorphine were selected to breed the APO-SUS rat strain. Apomorphine injection and behavioral selection were performed with the first 15 generations of APO-SUS rats. From the 15^th^ generation, APO-SUS rats displayed behavioral alterations without the apomorphine injection. In this study, a total of 94 male and female APO-SUS offspring and 103 normal Wistar rat offspring were used (n=197). Breeding of the litters consisted of placing a male rat in the cage of a female rat and leave them until a vaginal plug was found (considered as day 1 of gestation). Pregnant rats were housed individually. The offspring were weaned on postnatal day (PND) 21 and randomly housed in groups of 2-5 rats of the same sex and group. All rats had food and water available *ad libitum* and were housed in IVC cages (Optirat GenII IVC rack system) in humidity-controlled (55-60%) and thermo-regulated (21±2^o^C) rooms, with a reversed 12:12-h day/night cycle (lights off at 7 a.m.).

### 2.2. Experimental Design

Maternal immune activation (MIA) was induced by injecting pregnant Wistar rats (n=3) and APO-SUS rats (n=3) with the viral mimic poly I:C potassium salt on gestational day 15 (GD15). Control Wistar rats (n=4) and APO-SUS rats (n=5) were injected with saline. Male and female offspring were randomly divided into four groups: (1) offspring from Wistar mothers injected with saline (control), (2) offspring from Wistar mothers injected with poly I:C (MIA), (3) offspring from APO-SUS mothers injected with saline (APO-SUS), and (4) offspring from APO-SUS mothers injected with poly I:C (APO-SUS+MIA). Heart rate variability (PND7 and 14), USVs (PND7 and 14), and anticipatory locomotor activity (adolescence and adulthood) were measured in male and female rats. Adolescence refers to PND31-40 and adulthood to PND56-66. Different groups of male and female rats were used for the perinatal experiments (PND7 and 14), the experiments during adolescence, and the experiments during adulthood (i.e., 12 groups in total: 4 groups for each of the 3 experimental periods).

### 2.3. Maternal Immune Activation

Poly I:C (Sigma-Aldrich, Auckland, New Zealand, product number P1530, CAS number 42414-50-0) was stored at -18°C until solutions were prepared freshly in the morning of the day of administration. The poly I:C powder was dissolved in saline at a concentration of 5mg/ml. To allow reannealing of the double-stranded RNA structure, the solution was heated up to 65°C and maintained at this temperature for 5 minutes before cooling down to room temperature (22°C). On gestational day 15, pregnant rats were subcutaneously injected with saline or poly I:C solution at a dose of 5 mg per kg of body weight. Maternal weight and body temperature (measured on the flank with an infrared thermometer, Pro’s Kit, MT-4612) were measured just before the injection of poly I:C or saline and 2 and 24h post injection and used as an indicator of an immune response to the injection. This data is missing for 1 control Wistar and 1 APO-SUS rat injected with saline. Rats were placed back in their cage and left undisturbed for the remainder of their pregnancy. A checklist with the methodological details of the MIA model can be found in the supplemental materials (Kentner et al., 2019). To reduce the litter effect, one to three male and female offspring per litter were used for each experimental group.

### 2.4. Electrocardiographic Data acquisition – Heart rate variability

Electrocardiograms (ECG) were made on PND7 and 14 using the ECGenie apparatus. Each pup was brought to the experimental room and placed alone in a small (2*5 cm size) rectangular ECG holder (with controlled temperature to keep the pup warm and reduce stress) without bedding. Heart beats were measured through the paws using dedicated recording pads. The rat was left undisturbed for 8 minutes before being brought back to its home cage. The same rats were used on PND 7 and 14. LabChart 8 software was used to quantify parameters involved in heart rate variability (HRV). We also calculated the average heart rate. We measured the high frequency (using Fast Fourier transformation) and the root mean square of successive heartbeat internal differences (RMSSD), as they have been suggested to primarily measure the influence of the parasympathetic nervous system on HRV. We also measured the low frequency as it reflects a mixture of parasympathetic and sympathetic contributions with greater sympathetic sensitivity. The low frequency/high frequency (LF/HF) ratio was calculated to assess a possible shift in dominance between the sympathetic and parasympathetic nervous system (Kidwell and Ellenbroek, 2018; Rajendra Acharya et al., 2006).

### 2.5. Ultrasonic vocalizations (USVs)

USV were measured on PND7 and PND14 in male and female rats as an index of social communication and emotional state. During the 8-minute ECG periods, USVs were recorded with an ultrasound sensitive microphone (ultraMic 250 kHz) placed 2-3cm above the test cage and connected to a computer using Audacity software with a sampling rate of 19200 Hz. DeepSqueak software was used to manually analyze the USVs by determining the total number of calls, average length of the calls (s), total length or the calls (s) and principal frequency (Hz).

### 2.6. Anticipatory Locomotor activity

The anticipatory locomotor experiment was performed during adolescence (PND 30-40) and with a separate cohort of animals in adulthood (PND56-66) to measure anticipatory apathy in rats. The locomotor activity chambers (Med Associates Inc., USA; model ENV-515) were plexiglass square shape boxes (42*42*30 cm) equipped with two banks of eight photoelectric infrared cells (each 2.5 cm apart) on each of the internal walls of the chamber and at a height of 2.5 cm and 5 cm.

The experiment lasted 10 consecutive days and was divided into three parts: the habituation and learning phase (days 1-4 and 7-8), the pre-anticipation phase (days 5-6), and the anticipation phase (days 9-10). Days 1-4 are considered as habituation to the arena and days 7-8 as habituation to anticipatory learning. From days 1 to 5, rats were placed in the activity chambers and allowed to freely move for 35 minutes. Virkon S disinfectant was used to clean the chambers after each test. At the end of days 1-5, two Froot Loops (sugary cereals) per rat were places in their home cage to expose them to the reward. The experiments on days 6-10 were similar to those on days 1-5, but 3 Froot Loops were placed in the chambers after 25 minutes. The first 25 minutes (i.e., before the Froot Loops reward was placed in the chambers) of the pre-anticipation and anticipation phases were compared within groups using the distance traveled and the number of rearings to determine possible changes in anticipatory behavior. Within group comparison of the locomotion and number of rearings between the two phases were used as indicators of anticipatory pleasure. The last 10 minutes (after the Froot Loops were given) were not included in the analysis.

### 2.7. Western Blot Analysis

**Sample Preparation**

Frozen brain samples were homogenized in radioimmunoprecipitation assay (RIPA) buffer containing Halt™ Protease Inhibitor Cocktail (EDTA-Free; ThermoFisher Scientific, 87785) at a ratio of 2000 µL of buffer per gram of tissue. Homogenization was performed using a SONOPLUS Mini20 Ultrasonic Homogenizer (Bandelin Electronic) at 3 Watts for 30 seconds, employing 5-second bursts with 10-second intervals, while samples were kept on ice. Following homogenization, samples were centrifuged at 14,000 × g for 15 minutes at 4 °C, and the supernatants (lysates) were stored at -80 °C until further use.

**Protein Quantification**

Total protein concentrations in each sample were determined using the Pierce™ BCA Protein Assay Kit (ThermoFisher Scientific, Q23225). Protein standards and samples (25 µL each) were dispensed into a microplate. A working reagent (200 µL) was added to each well, and the plate was mixed for 30 seconds on a plate shaker. The plate was then incubated at 37 °C for 30 minutes. Absorbance was measured at 562 nm using a plate reader. Protein concentrations were determined by comparing sample absorbance values to a standard curve generated from known protein concentrations.

**Gel Electrophoresis and Membrane Transfer**

Protein samples were diluted in RIPA buffer to a final concentration of 4000 µg/mL, further diluted 1:1 in Laemmli buffer (containing 1:20 β-mercaptoethanol), and denatured at 95 °C for 5 minutes. Proteins were separated by sodium dodecyl sulfate-polyacrylamide gel electrophoresis (SDS-PAGE) using Mini-PROTEAN® TGX™ Precast Gels (Bio-Rad, 456-1085). Precision Plus Protein dual color standard (Bio-Rad, 161-0374) was loaded in the first lane. Twenty micrograms (10 µL) of each sample were loaded per lane, and separation was performed at 120 V for 90 minutes in SDS-PAGE running buffer using the Mini-PROTEAN® Tetra Vertical Electrophoresis Cell (Bio-Rad, 165-8004).

Subsequent to electrophoresis, proteins were transferred to Immobilon-P Transfer polyvinylidene difluoride (PVDF) membranes (Millipore, IPVH00010). Membranes were activated in methanol before assembling the transfer sandwich (grey side of cassette, fiber pad, filter paper, gel, PVDF membrane, filter paper, fiber pad, clear side of cassette). Transfer was conducted at 300 mA for 2 hours in cold Western blot transfer buffer using the Mini Trans-Blot® transfer tank (Bio-Rad, 170-3930), with the tank kept on ice.

**Protein Detection**

Post-transfer, membranes were washed once with tris-buffered saline with 0.1% Tween-20 (TBST) and blocked in 5% skim milk powder in TBST for 1 hour at room temperature. Following blocking, membranes were washed with TBST and incubated with primary antibodies overnight at 4 °C. The primary antibodies used were anti-PSD-95 (rabbit, 1:1000; Abcam, ab18258) and anti-Synaptophysin (rabbit, 1:2000; Abcam, ab52636), diluted in 1% skim milk powder in TBST. The following day, membranes were washed three times with TBST and incubated with the appropriate secondary antibodies for 1 hour at room temperature in the absence of light. The secondary antibodies used were AlexaFluor™ 488-conjugated anti-rabbit (1:5000; Abcam, ab150077) and AlexaFluor™ 488-conjugated anti-mouse (1:5000; Abcam, ab150113). Membranes were then washed three times with TBST and scanned using a Typhoon FLA 9000 scanner (GE Healthcare Bio-Sciences) at 500 V with the 473 nm laser and FITC filter.

To re-probe membranes for Alpha-tubulin as a loading control, membranes were stripped using a stringent stripping buffer consisting of 0.5 M Tris HCl (pH 6.8), 10% sodium dodecyl sulfate (SDS), 2-mercaptoethanol, and deionized water. The stripping buffer recipe included 12.5 mL of 0.5 M Tris HCl (pH 6.8), 20 mL of 10% SDS, 0.8 mL of 2-mercaptoethanol, and 67.5 mL of deionized water. Membranes were incubated in this buffer for 30 minutes at room temperature, washed six times with TBST, and re-scanned to ensure removal of previous antibody signals. Membranes were then blocked again in 5% skim milk powder in TBST for 1 hour, washed, and incubated with primary antibody against Alpha-tubulin (mouse, 1:5000; Abcam, ab7291) overnight at 4 °C. Following incubation with the primary antibody, membranes were washed, incubated with the secondary antibody, and scanned as described above.

**Data Analysis**

Western blot images were analyzed using Fiji software (ImageJ). Regions of interest (ROIs) were manually drawn around the bands corresponding to PSD-95 and Synaptophysin, as well as the background for each membrane. Background intensity was subtracted from the band intensity, and the band intensity values were normalized to the median intensity of the corresponding protein on each gel. Quantitative data were expressed as the corrected band intensity relative to the area of the ROI, with PSD-95 having an average area of 0.119 and Synaptophysin an average area of 0.127. This normalization ensures accurate comparisons across different gels and experimental conditions.

**Supplementary Table 1: Wald Chi-square (W), degrees of freedom (df) and p-values for main effects and interactions from the GEE analysis of the HRV, UVs, anticipatory locomotor tests and western blots**. HRV= Heart rate variability, UVS = ultrasonic vocalizations, ns = non-significant. Non-significant main effects and interactions are not shown.

| **Parameter** | **Value** | **Wald Chi-square/ F-value** | **Degrees of freedom** | **P value** |
| --- | --- | --- | --- | --- |
| USVs number of calls male | Main effect Genotype | 14.7 | 1 | **<0.001** |
|  | Main effect MIA | 1.170 | 1 | 0.279 |
|  | Main effect Time | 12.3 | 1 | **<0.001** |
|  | Interaction Genotype*MIA | 1.819 | 1 | 0.177 |
|  | Interaction Genotype*Time | 0.592 | 1 | 0.442 |
|  | Interaction MIA*Time | 0.344 | 1 | 0.558 |
|  | Interaction Time*MIA*Genotype | 2.004 | 1 | 0.157 |
| USVs Average length calls male | Main effect Genotype | 10.7 | 1 | **<0.001** |
|  | Main effect MIA | 0.800 | 1 | 0.371 |
|  | Main effect Time | 14.0 | 1 | **<0.001** |
|  | Interaction Genotype*MIA | 1.083 | 1 | 0.298 |
|  | Interaction Genotype*Time | 5.4 | 1 | 0.020 |
|  | Interaction MIA*Time | 0.672 | 1 | 0.413 |
|  | Interaction Time*MIA*Genotype | 4.427 | 1 | **0.035** |
| USVs Total length calls male | Main effect Genotype | 11.9 | 1 | **<0.001** |
|  | Main effect MIA | 0.065 | 1 | 0.799 |
|  | Main effect Time | 0.042 | 1 | 0.837 |
|  | Interaction Genotype*MIA | 0.598 | 1 | 0.480 |
|  | Interaction Genotype*Time | 0.695 | 1 | 0.404 |
|  | Interaction MIA*Time | 0.145 | 1 | 0.703 |
|  | Interaction Time*MIA*Genotype | 0.052 | 1 | 0.819 |
| USVs Principal frequency male | Main effect Genotype | 24.9 | 1 | **<0.001** |
|  | Main effect MIA | 2.049 | 1 | 0.152 |
|  | Main effect Time | 10.6 | 1 | **<0.001** |
|  | Interaction Genotype*MIA | 0.205 | 1 | 0.651 |
|  | Interaction Genotype*Time | 12.4 | 1 | **<0.001** |
|  | Interaction MIA*Time | 0.565 | 1 | 0.452 |
|  | Interaction Time*MIA*Genotype | 0.170 | 1 | 0.680 |
| USVs number of calls female | Main effect Genotype | 9.7 | 1 | **0.002** |
|  | Main effect MIA | 0.171 | 1 | 0.679 |
|  | Main effect Time | 18.1 | 1 | **<0.001** |
|  | Interaction Genotype*MIA | 0.002 | 1 | 0.969 |
|  | Interaction Genotype*Time | 0.097 | 1 | 0.755 |
|  | Interaction MIA*Time | 2.529 | 1 | 0.112 |
|  | Interaction Time*MIA*Genotype | 1.712 | 1 | 0.191 |
| USVs Average length calls female | Main effect Genotype | 25.0 | 1 | **<0.001** |
|  | Main effect MIA | 1.691 | 1 | 0.193 |
|  | Main effect Time | 5.4 | 1 | **0.021** |
|  | Interaction Genotype*MIA | 0.504 | 1 | 0.478 |
|  | Interaction Genotype*Time | 13.1 | 1 | **<0.001** |
|  | Interaction MIA*Time | 5.4 | 1 | **0.020** |
|  | Interaction Time*MIA*Genotype | 0.224 | 1 | 0.636 |
| USVs Total length calls female | Main effect Genotype | 23.7 | 1 | **<0.001** |
|  | Main effect MIA | 1.790 | 1 | 0.181 |
|  | Main effect Time | 3.404 | 1 | 0.065 |
|  | Interaction Genotype*MIA | 1.243 | 1 | 0.265 |
|  | Interaction Genotype*Time | 0.658 | 1 | 0.417 |
|  | Interaction MIA*Time | 2.258 | 1 | 0.071 |
|  | Interaction Time*MIA*Genotype | 1.555 | 1 | 0.212 |
| USVs Principal frequency female | Main effect Genotype | 18.0 | 1 | **<0.001** |
|  | Main effect MIA | 0.031 | 1 | 0.859 |
|  | Main effect Time | 5.1 | 1 | **0.024** |
|  | Interaction Genotype*MIA | 0.670 | 1 | 0.413 |
|  | Interaction Genotype*Time | 5.2 | 1 | **0.023** |
|  | Interaction MIA*Time | 2.461 | 1 | 0.117 |
|  | Interaction Time*MIA*Genotype | 0.458 | 1 | 0.499 |
| HRV Heart rate male | Main effect Genotype | 8.0 | 1 | **0.005** |
|  | Main effect MIA | 1.779 | 1 | 0.182 |
|  | Main effect Time | 497 | 1 | **<0.001** |
|  | Interaction Genotype*MIA | 5.5 | 1 | **0.019** |
|  | Interaction Genotype*Time | 0.954 | 1 | 0.351 |
|  | Interaction MIA*Time | 0.869 | 1 | 0.329 |
|  | Interaction Time*MIA*Genotype | 0.094 | 1 | 0.759 |
| HRV Low frequency male | Main effect Genotype | 0.064 | 1 | 0.800 |
|  | Main effect MIA | 0.721 | 1 | 0.396 |
|  | Main effect Time | 18.4 | 1 | **<0.001** |
|  | Interaction Genotype*MIA | 0.390 | 1 | 0.532 |
|  | Interaction Genotype*Time | 0.018 | 1 | 0.893 |
|  | Interaction MIA*Time | 0.617 | 1 | 0.432 |
|  | Interaction Time*MIA*Genotype | 0.247 | 1 | 0.619 |
| HRV High frequency male | Main effect Genotype | 4.8 | 1 | **0.028** |
|  | Main effect MIA | 2.343 | 1 | 0.126 |
|  | Main effect Time | 2.088 | 1 | 0.148 |
|  | Interaction Genotype*MIA | 0.896 | 1 | 0.372 |
|  | Interaction Genotype*Time | 0.002 | 1 | 0.961 |
|  | Interaction MIA*Time | 1.063 | 1 | 0.301 |
|  | Interaction Time*MIA*Genotype | 0.068 | 1 | 0.794 |
| HRV LF/HF ratio male | Main effect Genotype | 0.161 | 1 | 0.689 |
|  | Main effect MIA | 0.534 | 1 | 0.465 |
|  | Main effect Time | 11.1 | 1 | **<0.001** |
|  | Interaction Genotype*MIA | 0.710 | 1 | 0.399 |
|  | Interaction Genotype*Time | 0.009 | 1 | 0.924 |
|  | Interaction MIA*Time | 1.996 | 1 | 0.158 |
|  | Interaction Time*MIA*Genotype | 0.101 | 1 | 0.750 |
| HRV RMSSD male | Main effect Genotype | 0.465 | 1 | 0.495 |
|  | Main effect MIA | 1.518 | 1 | 0.218 |
|  | Main effect Time | 0.079 | 1 | 0.779 |
|  | Interaction Genotype*MIA | 0.234 | 1 | 0.629 |
|  | Interaction Genotype*Time | 0.169 | 1 | 0.681 |
|  | Interaction MIA*Time | 1.953 | 1 | 0.162 |
|  | Interaction Time*MIA*Genotype | 0.331 | 1 | 0.565 |
| HRV Heart rate female | Main effect Genotype | 6.4 | 1 | **0.010** |
|  | Main effect MIA | 0.571 | 1 | 0.450 |
|  | Main effect Time | 607.1 | 1 | **<0.001** |
|  | Interaction Genotype*MIA | 0.429 | 1 | 0.513 |
|  | Interaction Genotype*Time | 6.1 | 1 | **0.010** |
|  | Interaction MIA*Time | 0.115 | 1 | 0.734 |
|  | Interaction Time*MIA*Genotype | 2.701 | 1 | 0.100 |
| HRV Low frequency female | Main effect Genotype | 3.355 | 1 | 0.067 |
|  | Main effect MIA | 1.886 | 1 | 0.170 |
|  | Main effect Time | 12.5 | 1 | **<0.001** |
|  | Interaction Genotype*MIA | 3.349 | 1 | 0.067 |
|  | Interaction Genotype*Time | 8.2 | 1 | **0.004** |
|  | Interaction MIA*Time | 1.496 | 1 | 0.221 |
|  | Interaction Time*MIA*Genotype | 0.855 | 1 | 0.328 |
| HRV High frequency female | Main effect Genotype | 0.476 | 1 | 0.490 |
|  | Main effect MIA | 0.555 | 1 | 0.456 |
|  | Main effect Time | 0.433 | 1 | 0.511 |
|  | Interaction Genotype*MIA | 4.1 | 1 | **0.004** |
|  | Interaction Genotype*Time | 1.532 | 1 | 0.216 |
|  | Interaction MIA*Time | 1.197 | 1 | 0.274 |
|  | Interaction Time*MIA*Genotype | 2.227 | 1 | 0.136 |
| HRV LF/HF ratio female | Main effect Genotype | 6.1 | 1 | **0.014** |
|  | Main effect MIA | 1.234 | 1 | 0.267 |
|  | Main effect Time | 7.3 | 1 | **0.007** |
|  | Interaction Genotype*MIA | 0.213 | 1 | 0.644 |
|  | Interaction Genotype*Time | 9.9 | 1 | **0.002** |
|  | Interaction MIA*Time | 2.310 | 1 | 0.129 |
|  | Interaction Time*MIA*Genotype | 1.658 | 1 | 0.198 |
| HRV RMSSD female | Main effect Genotype | 4.0 | 1 | **0.047** |
|  | Main effect MIA | 0.417 | 1 | 0.518 |
|  | Main effect Time | 0.368 | 1 | 0.544 |
|  | Interaction Genotype*MIA | 6.2 | 1 | **0.013** |
|  | Interaction Genotype*Time | 1.451 | 1 | 0.228 |
|  | Interaction MIA*Time | 0.231 | 1 | 0.630 |
|  | Interaction Time*MIA*Genotype | 0.006 | 1 | 0.937 |
| Adolescence Distance travelled male | Main effect Genotype | 7.5 | 1 | **0.006** |
|  | Main effect MIA | 3.256 | 1 | 0.071 |
|  | Main effect Time | 17.5 | 1 | **<0.001** |
|  | Interaction Genotype*MIA | 0.238 | 1 | 0.625 |
|  | Interaction Genotype*Time | 0.315 | 1 | 0.574 |
|  | Interaction MIA*Time | 0.447 | 1 | 0.504 |
|  | Interaction Time*MIA*Genotype | 2.172 | 1 | 0.141 |
| Adolescence Rearing male | Main effect Genotype | 3.9 | 1 | **0.047** |
|  | Main effect MIA | 15.6 | 1 | **0.010** |
|  | Main effect Time | 0.419 | 1 | 0.517 |
|  | Interaction Genotype*MIA | 0.973 | 1 | 0.324 |
|  | Interaction Genotype*Time | 4.6 | 1 | **0.032** |
|  | Interaction MIA*Time | 0.201 | 1 | 0.654 |
|  | Interaction Time*MIA*Genotype | 6.471 | 1 | **0.011** |
| Adulthood Distance travelled male | Main effect Genotype | 7.4 | 1 | **0.007** |
|  | Main effect MIA | 0.139 | 1 | 0.709 |
|  | Main effect Time | 1.154 | 1 | 0.228 |
|  | Interaction Genotype*MIA | 0.021 | 1 | 0.886 |
|  | Interaction Genotype*Time | 4.8 | 1 | **0.029** |
|  | Interaction MIA*Time | 1.949 | 1 | 0.163 |
|  | Interaction Time*MIA*Genotype | 0.002 | 1 | 0.961 |
| Adulthood Rearing male | Main effect Genotype | 6.5 | 1 | **0.011** |
|  | Main effect MIA | 0.453 | 1 | 0.501 |
|  | Main effect Time | 9.8 | 1 | **0.002** |
|  | Interaction Genotype*MIA | 0.567 | 1 | 0.452 |
|  | Interaction Genotype*Time | 1.667 | 1 | 0.197 |
|  | Interaction MIA*Time | 0.093 | 1 | 0.761 |
|  | Interaction Time*MIA*Genotype | 0.001 | 1 | 0.980 |
| Adolescence Distance travelled female | Main effect Genotype | 4.6 | 1 | **0.032** |
|  | Main effect MIA | 0.709 | 1 | 0.400 |
|  | Main effect Time | 26 | 1 | **<0.001** |
|  | Interaction Genotype*MIA | 2.511 | 1 | 0.113 |
|  | Interaction Genotype*Time | 0.590 | 1 | 0.442 |
|  | Interaction MIA*Time | 14.2 | 1 | **<0.001** |
|  | Interaction Time*MIA*Genotype | 5.631 | 1 | **0.018** |
| Adolescence Rearing female | Main effect Genotype | 11.6 | 1 | **<0.001** |
|  | Main effect MIA | 3.989 | 1 | **0.046** |
|  | Main effect Time | 11.2 | 1 | **<0.001** |
|  | Interaction Genotype*MIA | 2.555 | 1 | 0.110 |
|  | Interaction Genotype*Time | 0.732 | 1 | 0.392 |
|  | Interaction MIA*Time | 0.001 | 1 | **<0.001** |
|  | Interaction Time*MIA*Genotype | 0.790 | 1 | 0.374 |
| Adulthood Distance travelled female | Main effect Genotype | 1.204 | 1 | 0.272 |
|  | Main effect MIA | 0.565 | 1 | 0.452 |
|  | Main effect Time | 3.053 | 1 | 0.081 |
|  | Interaction Genotype*MIA | 2.264 | 1 | 0.125 |
|  | Interaction Genotype*Time | 1.803 | 1 | 0.179 |
|  | Interaction MIA*Time | 15.526 | 1 | **<0.001** |
|  | Interaction Time*MIA*Genotype | 1.934 | 1 | 0.164 |
| Adulthood Rearing female | Main effect Genotype | 1.626 | 1 | 0.202 |
|  | Main effect MIA | 5.4 | 1 | **0.020** |
|  | Main effect Time | 9.8 | 1 | **0.002** |
|  | Interaction Genotype*MIA | 1.680 | 1 | 0.195 |
|  | Interaction Genotype*Time | 1.869 | 1 | 0.172 |
|  | Interaction MIA*Time | 6.7 | 1 | **0.010** |
|  | Interaction Time*MIA*Genotype | 0.590 | 1 | 0.442 |
| Hippocampus Adolescence Synaptophysin male | Main effect Genotype | F (1, 20) = 3.387 | 1 | 0.081 |
|  | Main effect MIA | F (1, 20) = 5.600 | 1 | **0.028** |
|  | Interaction Genotype*MIA | F (1, 20) = 0.844 | 1 | 0.369 |
| Hippocampus Adulthood Synaptophysin male | Main effect Genotype | F (1, 20) = 19.60 | 1 | **0.0003** |
|  | Main effect MIA | F (1, 20) = 19.35 | 1 | **0.0003** |
|  | Interaction Genotype*MIA | F (1, 20) = 8.485 | 1 | **0.009** |
| Hippocampus Adolescence Synaptophysin female | Main effect Genotype | F (1, 20) = 4.839 | 1 | **0.040** |
|  | Main effect MIA | F (1, 20) = 0.574 | 1 | 0.457 |
|  | Interaction Genotype*MIA | F (1, 20) = 7.989 | 1 | **0.010** |
| Hippocampus Adulthood Synaptophysin female | Main effect Genotype | F (1, 20) = 0.728 | 1 | 0.404 |
|  | Main effect MIA | F (1, 20) = 0.859 | 1 | 0.365 |
|  | Interaction Genotype*MIA | F (1, 20) = 7.330 | 1 | **0.014** |
| Hippocampus Adolescence  PSD95 male | Main effect Genotype | F (1, 20) = 8.079 | 1 | **0.010** |
|  | Main effect MIA | F (1, 20) = 0.021 | 1 | 0.885 |
|  | Interaction Genotype*MIA | F (1, 20) = 8.079 | 1 | 0.774 |
| Hippocampus Adulthood  PSD95 male | Main effect Genotype | F (1, 19) = 2.499 | 1 | 0.130 |
|  | Main effect MIA | F (1, 19) = 1.138 | 1 | 0.230 |
|  | Interaction Genotype*MIA | F (1, 19) = 1.709 | 1 | 0.207 |
| Hippocampus Adolescence  PSD95 female | Main effect Genotype | F (1, 18) = 2.311 | 1 | 0.146 |
|  | Main effect MIA | F (1, 18) = 0.145 | 1 | 0.707 |
|  | Interaction Genotype*MIA | F (1, 18) = 0.009 | 1 | 0.926 |
| Hippocampus Adulthood  PSD95 female | Main effect Genotype | F (1, 20) = 0.206 |  | 0.655 |
|  | Main effect MIA | F (1, 20) = 7.921 | 1 | **0.011** |
|  | Interaction Genotype*MIA | F (1, 20) = 2.121 |  | 0.161 |
| Frontal cortex Adolescence Synaptophysin male | Main effect Genotype | F (1, 19) = 10.38 | 1 | **0.0045** |
|  | Main effect MIA | F (1, 19) = 14.01 | 1 | **0.0014** |
|  | Interaction Genotype*MIA | F (1, 19) = 17.04 | 1 | **0.0006** |
| Frontal cortex Adolescence Synaptophysin male | Interaction Genotype*MIA | F (1, 20) = 0.038 | 1 | 0.846 |
|  | Main effect Genotype | F (1, 20) = 0.374 | 1 | 0.547 |
|  | Main effect MIA | F (1, 20) = 0.322 | 1 | 0.577 |
| Frontal cortex Adolescence Synaptophysin female | Main effect Genotype | F (1, 20) = 6.989 | 1 | **0.016** |
|  | Main effect MIA | F (1, 20) = 8.833 | 1 | **0.008** |
|  | Interaction Genotype*MIA | F (1, 20) = 0.021 | 1 | 0.884 |
| Frontal cortex Adulthood Synaptophysin female | Main effect Genotype | F (1, 20) = 0.079 |  | 0.780 |
|  | Main effect MIA | F (1, 20) = 4.132 |  | 0.056 |
|  | Interaction Genotype*MIA | F (1, 20) = 14.22 | 1 | **0.0012** |
| Frontal cortex Adolescence  PSD95 male | Main effect Genotype | F (1, 19) = 0.751 | 1 | 0.464 |
|  | Main effect MIA | F (1, 19) = 8.409 | 1 | **0.009** |
|  | Interaction Genotype*MIA | F (1, 19) = 0.557 | 1 | 0.397 |
| Frontal cortex Adulthood  PSD95 male | Main effect Genotype | F (1, 20) = 1.365 | 1 | 0.256 |
|  | Main effect MIA | F (1, 20) = 1.010 | 1 | 0.327 |
|  | Interaction Genotype*MIA | F (1, 20) = 0.788 | 1 | 0.385 |
| Frontal cortex Adolescence  PSD95 female | Main effect Genotype | F (1, 20) = 0.070 | 1 | 0.793 |
|  | Main effect MIA | F (1, 20) = 2.260 | 1 | 0.148 |
|  | Interaction Genotype*MIA | F (1, 20) = 4.436 | 1 | **0.048** |
| Frontal cortex Adulthood  PSD95 female | Main effect Genotype | F (1, 20) = 0.171 | 1 | 0.683 |
|  | Main effect MIA | F (1, 20) = 6.546 | 1 | **0.019** |
|  | Interaction Genotype*MIA | F (1, 20) = 0.345 | 1 | 0.563 |

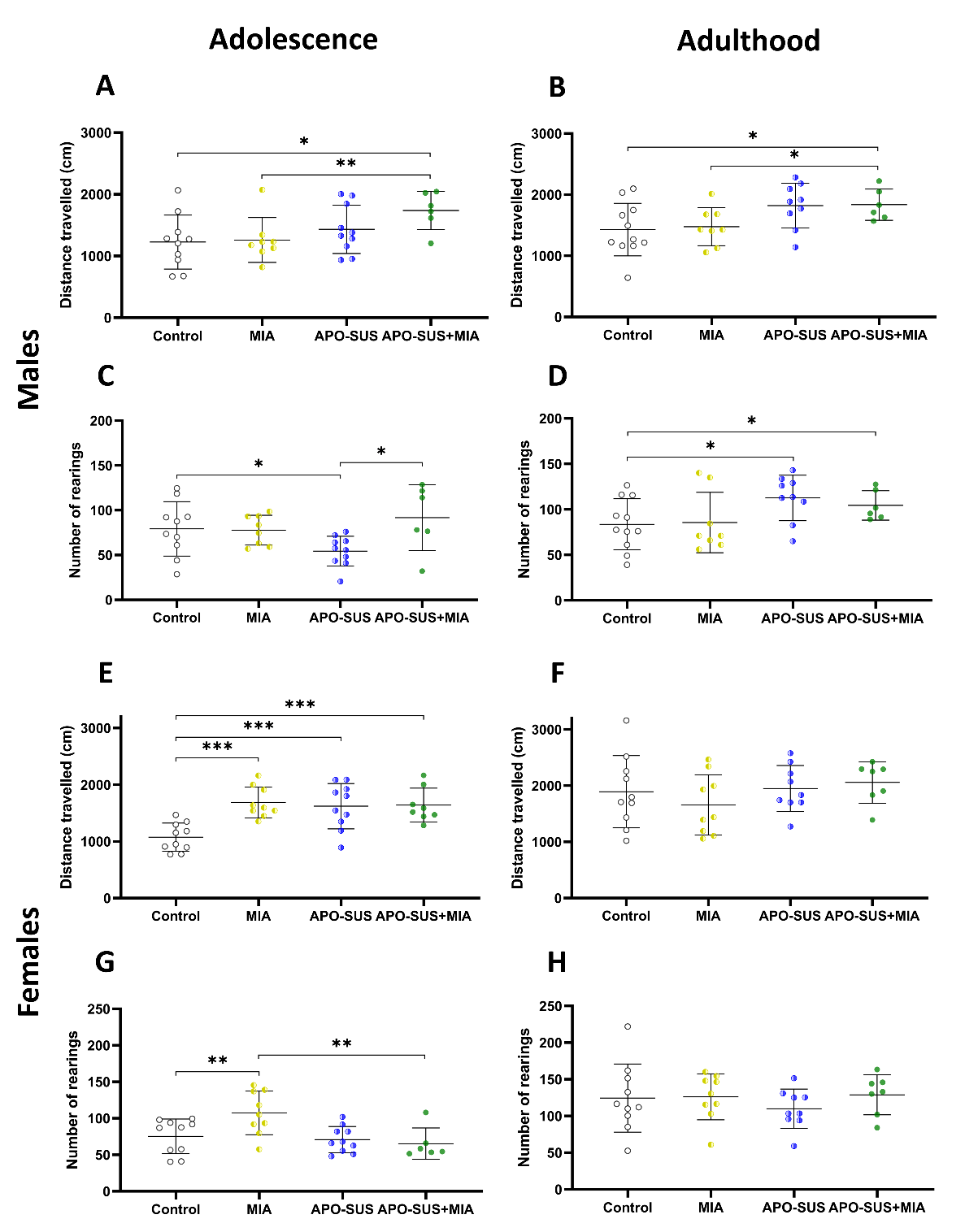

**Supplementary Figure 1. *Between group comparison of the locomotor and rearing activity in control, MIA, APO-SUS, and APO-SUS+MIA rats.*** *Distance traveled in males during adolescence (****A****.) and adulthood (****B****.). Number of rearings in males during adolescence (****C****.) and adulthood (****D****.). Distance traveled in females during adolescence (****E****.) and adulthood (****F****.). Number of rearings in females during adolescence (****G****.) and adulthood (****H****.). N=6-10 males and 7-10 females per group. Statistically significant differences between groups are indicated by asterisks: *p<0.05, **p<0.01, ***p<0.001. Data is presented as mean± SD. Significant differences between time points and between MIA and APO-SUS are not shown.*
